# Testing hypotheses about correlations between brain activation patterns

**DOI:** 10.64898/2026.03.21.713393

**Authors:** Jörn Diedrichsen, Xianglong Fu, Mahdiyar Shahbazi, Simon Bonner

## Abstract

Many functional magnetic resonance imaging (fMRI) studies conclude that two conditions engage “overlapping, yet partly distinct” patterns of activation. Yet, there is currently no commonly accepted method for determining the extent of this overlap. While correlations between activation patterns can serve as a measure of their correspondence, empirical correlations are strongly biased towards zero due to measurement noise, preventing their use in testing hypotheses about the actual degree of pattern correspondence. In this paper, we derive the maximum-likelihood estimate for the correlation of the true (noise-less) activation patterns and examine its behavior in the low signal-to-noise regime that is typical for fMRI studies. We show that although the maximum-likelihood estimate corrects for much of the influence of measurement noise, it is ultimately biased. We examine different ways of drawing inferences about the size of the underlying true correlations. We find that a subject-wise bootstrap on the maximum-likelihood group estimate performs best over the tested conditions. We extend the proposed method to test more general hypotheses about the representational geometry of activation patterns for more conditions, and highlight best practices, as well as common pitfalls and problems, in testing such hypotheses.

## 1 INTRODUCTION

Most functional magnetic resonance imaging (fMRI) studies compare brain activity across different experimental conditions. For example, one may ask whether the planning of a movement causes an activation pattern that is similar to executing the same movement. Often such studies conclude that two conditions elicit “overlapping, yet partly distinct” patterns of activation (e.g., Beffara et al., 2023; Feola et al., 2023; Kabulska et al., 2024; Pan et al., 2025; Guo et al., 2023). The exact amount of correspondence of the activation pattern is an important quantity here; it indicates to what degree the two conditions rely on common neural processes in that specific brain region. In turn, the amount of non-overlap indicates to what degree the two conditions engage separate processes.

Given the neuroscientific importance of the true degree of overlap, it is remarkable that there is no commonly accepted way of drawing statistical inferences about the degree of correspondence of spatial (i.e. multivariate) activation patterns. At first sight, the obvious solution to this problem is to calculate the correlation between the two activation patterns across voxels (the spatial measurement unit of fMRI). Correlations are clearly an adequate measure here, because when we judge the degree of overlap, we are not interested in the absolute size of the activation patterns. For example, planning a movement will elicit much weaker activity than executing it - yet the two conditions may still induce the same pattern of activation across voxels, causing the two activation patterns to be highly correlated.^1^ This would suggest that planning a movement activates exactly the same network as executing it.

The problem is that we do not have access to the true activation patterns, but we need to rely on noisy measurements. Let the vectors **x**^*\**^ and **y**^*\**^ denote the true activity values for the two conditions for *P* voxels. When running an fMRI experiments, we obtain *n*_*x*_ measurements for the first condition *{***x**_1_, .., **x**_*nx*_ *}*, and *n*_*y*_ measurements for the second condition *{***y**_1_, …, **y**_*ny*_*}*. These can then be averaged to obtain the average pattern estimates 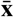 and 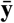 (Fig. 1a). The correlation between the two averaged patterns underestimates the true correlation (*ρ*) substantially, as it overestimates the variance of the true patterns 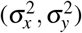. This bias can be substantial when we analyze unsmoothed single-subject fMRI data, where the variance of measurement noise, even after averaging, often outstrips the true patterns by orders of magnitude.

**Figure 1.**
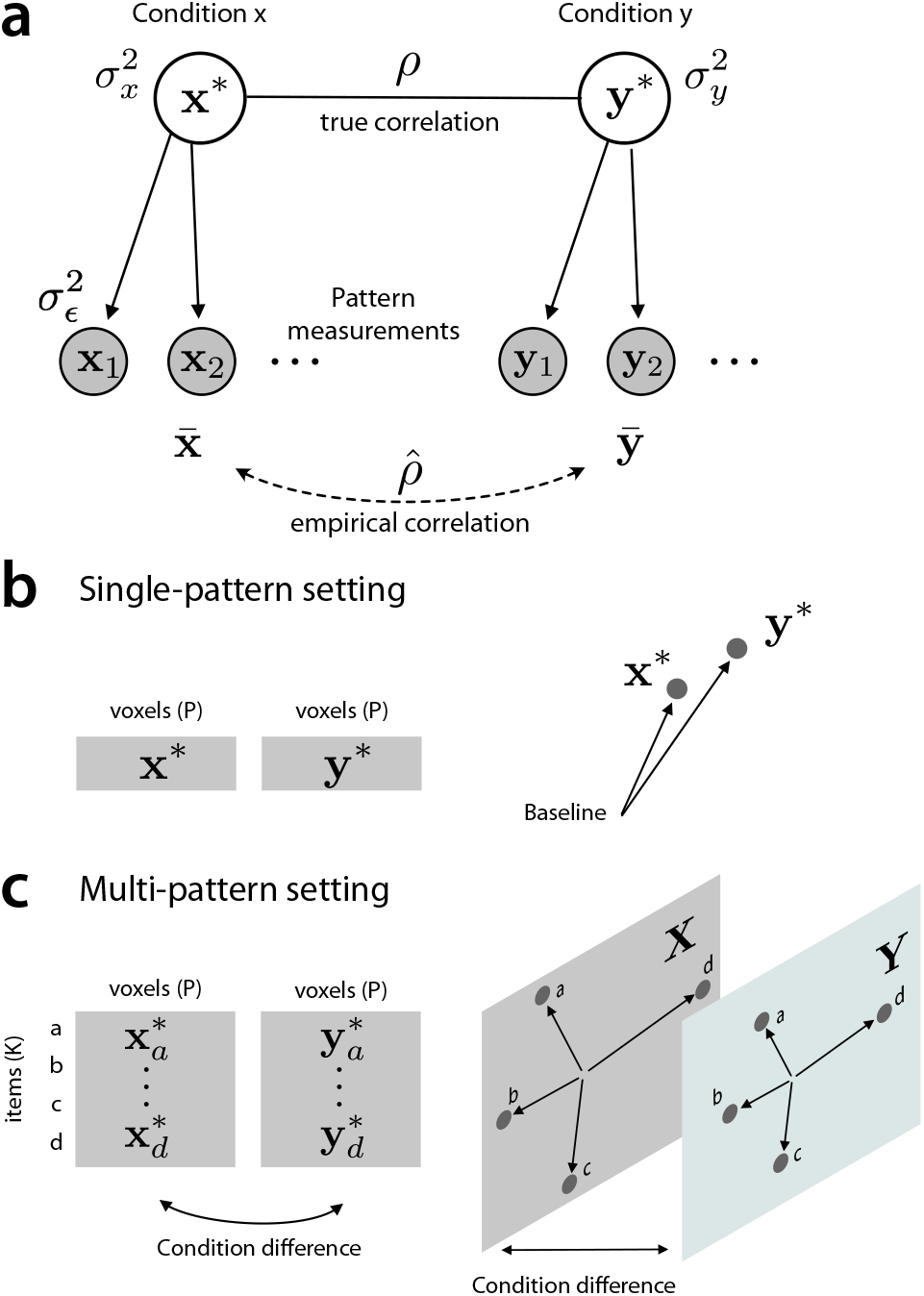
Problem statement. **(a)** A graphical model of the problem. The true activation patterns (**x**^*\**^ and **y**^*\**^) have spatial variance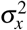 and 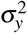 across *P* voxels. We obtain multiple measurements (gray circles) for each pattern with measurement noise 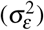. The correlation between the mean activation patterns 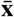 and 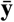 will underestimate the true correlation. **(b)** In the simplest case we are interested in the correlation / cosine similarity between the two patterns relative to a common baseline. The cosine similarity between two patterns is the cosine of the angle between the two *P*-dimensional vectors. **(c)** In the more advanced multi-pattern setting, we measure two or more items under each condition, and want to establish if differences between the items (or between each item and the condition mean) are parallel across conditions.

The existence of this bias has long been recognized in the statistical literature (Spearman, 1904), and a number of related techniques to correct for the bias have been proposed in various scientific fields (Beaton et al., 1979; Saccenti et al., 2020; Liu et al., 1978). Building on these ideas, the contribution of the current paper is four-fold. First, we introduce the maximum-likelihood estimator as a slightly more general version of the point estimators for a corrected correlations. Secondly, using simulations, we study the bias and stability of this estimator in very low functional signal-to-noise ratio (fSNR) regimes, as it is typical for unsmoothed single-subject fMRI data. Third, we test different methods of how to draw inferences on hypotheses involving these correlation coefficients. Specifically we consider the hypotheses that the correlation between two conditions is larger or smaller than a fixed value (one-sample inference), and the hypothesis that the correlation differs between two pairs of conditions or regions (paired-sample inference). Finally, we show that the proposed methods can be used to infer on correlations in two related scenarios, both of which occur quite often in fMRI studies.

In the simple single-pattern setting (Fig. 1b) there is only one true activation pattern per condition. Each of these (**x**^*\**^ and **y**^*\**^) is a *P*-dimensional vector, with each dimension reflecting the activity of a single voxel in that condition compared to rest (Fig. 1b). In this setting we are interested in the cosine similarity between the two patterns, the angle between the two vectors that connect each of the conditions to a common baseline.

In the more complex multi-pattern setting (Fig. 1c), we measure the activation patterns for multiple stimuli or items under each of the two conditions. In this problem, which often arises in multi-voxel pattern analysis fMRI studies (Kriegeskorte et al., 2006; Kriegeskorte, 2011; Norman et al., 2006), we are not interested in the correlation between the mean condition patterns relative to a common baseline, but rather whether the activation patterns that differentiate between different items are correlated across conditions. In essence we seek to establish the cosine of the angle between the vectors connecting the item-specific patterns across the two conditions conditions. For example, we may measure the activation patterns, while a subject either executes four different actions, or observes the same actions (Oosterhof et al., 2010; Gazzola and Keysers, 2009). If the differences in the item-specific activation patterns are correlated across the two conditions, then the region uses similar neuronal patterns to represent specific actions, even though the mean activity for execution or observing that action may be dramatically different. Traditionally the correspondence of representations has been addressed using cross-decoding approaches, training a classifier to distinguish the different items in one condition and then classifying the items in the other condition (Gallivan et al., 2013; Dinstein et al., 2008; Formisano et al., 2008; Harrison and Tong, 2009; Gallivan et al., 2011). This method, however, does not easily allow us to determine the exact degree of representational alignment. We show here that our methods can also be applied in the multi-pattern scenario to provide valid inferences on the degree of correspondence between two sets of activation patterns.

## 2 METHODS

### 2.1 Definitions

We are interested in the correlation or cosine similarity between the activation patterns related to two experimental conditions. Typically, these activation patterns are measured in *s* = 1, …, *S* subjects, each across *p* = 1, …, *P*^(*s*)^ voxels of a specific brain region. The true activity values for voxel *p* in subject *s* are denoted as 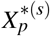 and 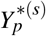. The true activation values differ across subjects and voxels, such that we consider them as a latent random variables. We assume that these true patterns have mean zero and variances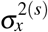 and 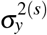, with the variance possibly different across subjects. For the next sections, we focus on the estimate of the correlation / cosine similarity from the dataset of a single subject and will therefore drop the the index (*s*) for notational simplicity. We will reintroduce the index again in the section on group estimates.

Suppose that we have *n*_*x*_ measurements for the first and *n*_*y*_ measurements for the second condition, with the total number of measurements being *N* = *n*_*x*_ + *n*_*y*_. The *i*^*th*^ measurements for the *p*^*th*^ voxel are modelled as

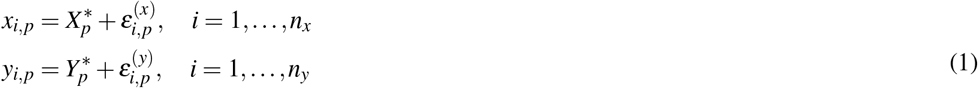

We assume that the measurement noise for *x*_*i,p*_ and *y*_*i,p*_ have mean 0 and the same variance 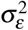. We also assume that the measurement noise is independent across conditions, voxels, and measurements (see section 3.3 for impact of dependence across voxels).

### 2.2 Correlations vs. cosine similarities

If we assume that the mean of the true patterns across voxels is zero, then the true variance and correlations between 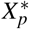 and 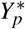 are:

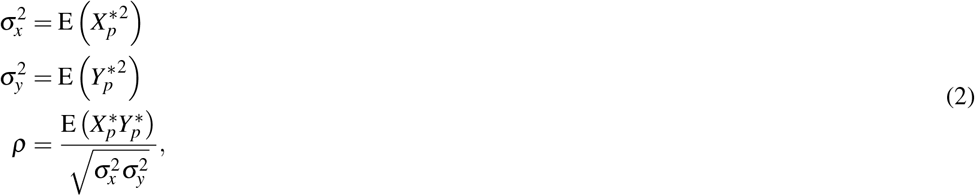

where all expectations (E) are across voxels. If the mean value across all voxels cannot be assumed to be zero, then we have two choices: First, we can subtract the mean of all patterns across voxels, such that the quantities in Eq. 2 are indeed variances and correlation. Alternatively, we can decide not to remove the mean. In this case, 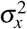 becomes the second moment of *X*^*\**^ and *ρ* the cosine-similarity between the true patterns. This choice is often preferable in multivariate fMRI analysis, as we want to take into account the mean activity across voxels when judging the similarity of patterns relative to a baseline condition (Walther et al., 2016). For the remainder of the paper we will refer to 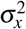 and 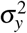 as variances and *ρ* as correlations, instead of calling them second moments and cosine similarities, knowing that our results pertain both situations.

### 2.3 Simple estimates

We can attempt to estimate *ρ* by averaging the pattern estimates across observations

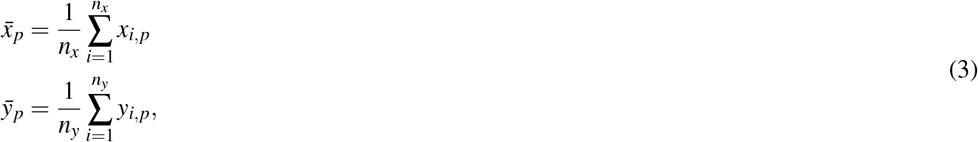

and then calculate the simple Pearson correlation between these average estimates

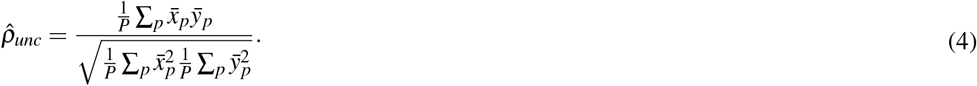

While the numerator is a unbiased estimator of E 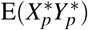, the denominator is not unbiased estimator of the signal variances. Rather the individual terms are positively biased by measurement noise:

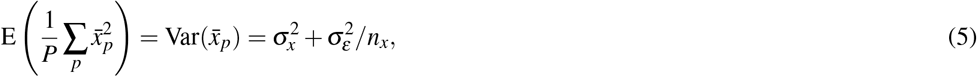

and similarly for 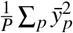. This positive bias causes the Pearson correlation of the means to underestimate the true correlation between two variables, a fact well recognized already by Spearman (1904).

### 2.4 Maximum likelihood estimates

To correct for the biasing influence of measurement noise, we can derive the maximum-likelihood estimate (MLE) for the variances and correlation under the assumption that both the signal and the measurement noise are normally distributed. This is the approach taken in pattern component modeling (PCM, Diedrichsen et al., 2011, 2018). In this framework, the concatinated true activity values at voxel *p* under both conditions, denoted by the **u**_*p*_, follow a bivariate normal random distribution with zero mean and variance-covariance matrix **G**(*θ*):

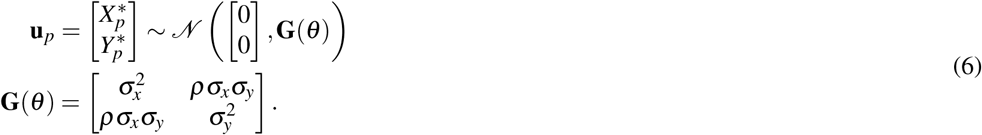

We also define **d**_*p*_ as the vector containing all *N* measurements for the *p*^*th*^ voxel, and **Z** as an *N×* 2 design matrix that indicates whether the *n*^*th*^ measurement belongs to the first or second condition. The measurement model (Eq. 1) for the data conditional on **u**_*p*_ can then be written as:

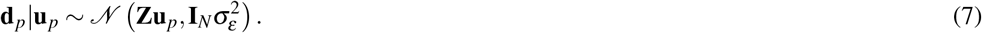

where 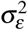 is the variance of the independent measurement noise. The marginal distribution of the data for voxel *p* is then:

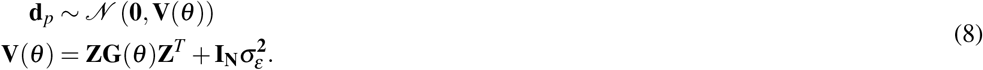

To derive the overall likelihood for a set of voxels, we concatenate the voxel measurements from a single subject into the *N× P* matrix (**D**). Assuming independence of the data across voxels (see section 3.3) the log-likelihood then can be written as:

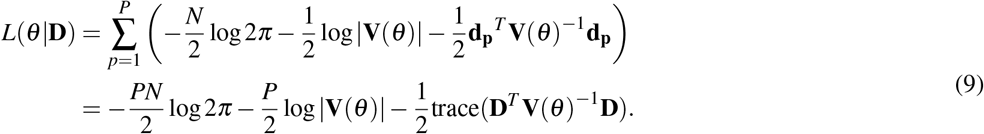

To maximize this quantity given the constraint that all *σ >* 0 and that *−*1 *< ρ <* 1, we apply a transform, such that our parameters *θ* are unbounded:

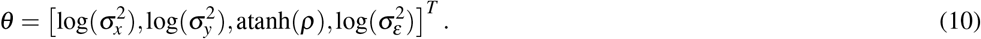

With this transform, the log-likelihood can be readily optimized using the first derivative in respect to each parameters (see Appendix 6.3 for details):

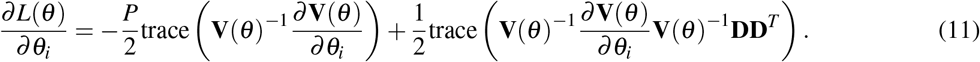

From *θ* we can then obtain the MLEs for the correlation correlation 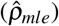 and the variances of the true patterns 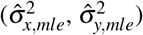, and measurement noise 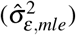 by inverting the transforms of Eq. 10.

If the MLE for the variance of the pattern for one of the conditions approaches 0, the predicted covariance of patterns does not depend on the value of *ρ* anymore (Eq. 6). Consequently, the likelihood function becomes flat in respect to *ρ*. When using the unbounded variable transform (Eq. 10), the MLE for the corelation will approach 1 for positive covariance between the conditions and *−*1 for negative covariance. In our analysis, we identify all estimates for which 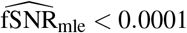 (see section 2.9) as having no signal. We show in the following that it is important to not exclude these estimates when drawing inferences.

### 2.5 Cross-block estimate

As long as the MLE of the parameters do not lie at the limits of the allowed range 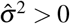 or 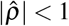, they can be derived by analytically matching the empirical variances and covariance to their expected values (method of moments). This approach has been suggested in a number of fields (Beaton et al., 1979; Rosner and Willett, 1988; Saccenti et al., 2020). For our specific measurement error model (assuming independent measures, with the same error variance for both conditions), the cross-block estimator (*cbe*) for the variances are:

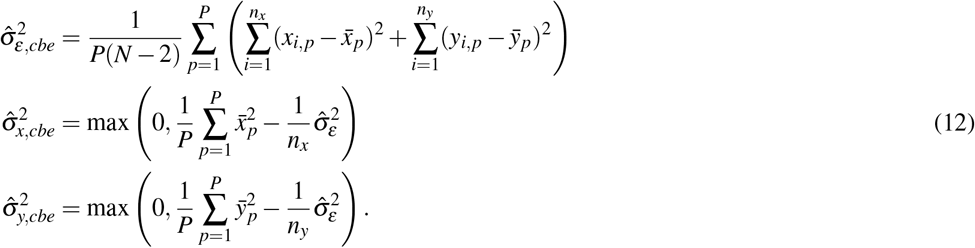

We can then compute the numerator

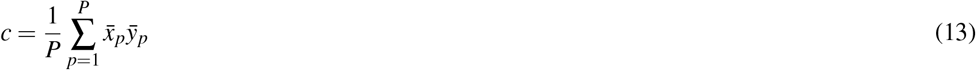

and denominator

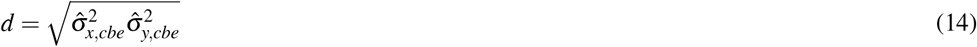

for the correlation estimator, which then needs to be constrained to lie in the interval from [*−*1, 1].

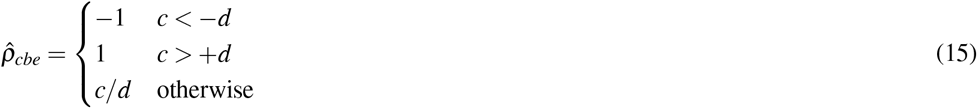

These quantities are easy to calculate and the correlation estimator 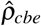 is usually very close to 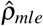. However when the maximum-likelihood estimate for the correlation lies on a boundary 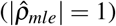 the variance estimators for these two approaches do not match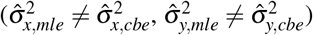. As we will see in the results, the cross-block estimators for the signal variances are often zero, even though the corresponding maximum-likelihood estimators are still positive. We will also show that the exclusion of cases with zero estimated variance can bias inferential procedures.

### 2.6 Multi-pattern setting, maximum-likelihood estimate

In the multi-pattern setting, we are measuring the activation pattern of *j* = 1…*K* items under the two conditions for a repeated number times *i* = 1…*n*_*x*_ and *i* = 1…*n*_*y*_. 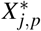 be the true activation pattern for item *j* and voxel *p*, and *x*_*i, j,p*_ denote the *i*^*th*^ observation thereof. We are interested in whether the differences between each item and the mean of each condition (across items) are parallel across the two conditions (Fig. 1c).

To derive the MLE, we define a vector of the activation values of the true condition means (averaged across items):

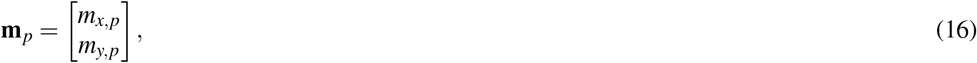

and **u**_*p*_, a 2*K*-long vector, consisting of the difference in true activation for each item from the corresponding condition means:

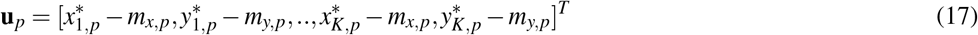

The data vector **d**_*p*_ then contains all measurements for that voxel (*x*_*i, j,p*_ and *y*_*i, j,p*_). The full measurement model can then be written as

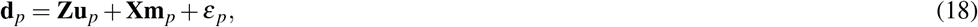

where **Z** is a *N×* 2*K* matrix that links each observation to the corresponding item / condition in the vector **u**_*p*_, **X** a *N×* 2 matrix that links each observation to the corresponding condition mean, and *ε* _*p*_ the vector of measurement noise terms, assumed to have independent normal distribution with zero mean and variance 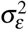. Under this model, the variance-covariance matrix of **u**_*p*_ is:

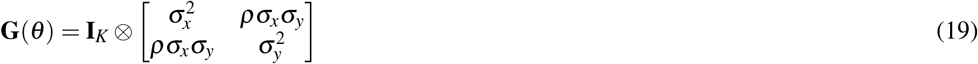

where ⊗denotes the Kronecker product.

When estimating the parameters we need to then take the removal of the condition mean into account. This is achieved by maximizing the Restricted Maximum Likelihood (Speed, 1997; Diedrichsen et al., 2018). For details see Appendix 6.4.

### 2.7 Multi-item setting, cross-block estimate

To define the corresponding cross-block estimate, we first subtract the mean of each condition (across items) from each measurement:

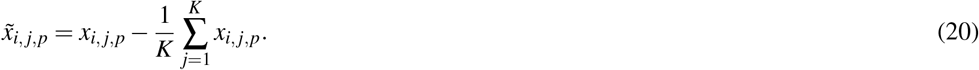

The mean of these values across repetitions 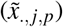 then gives us an estimate of the pattern by which a specific item differs from the condition mean:

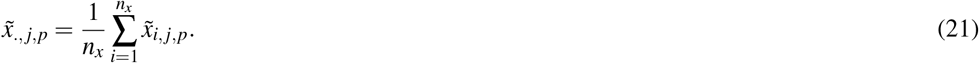

Using this estimate, the cross-block estimators of the variances are:

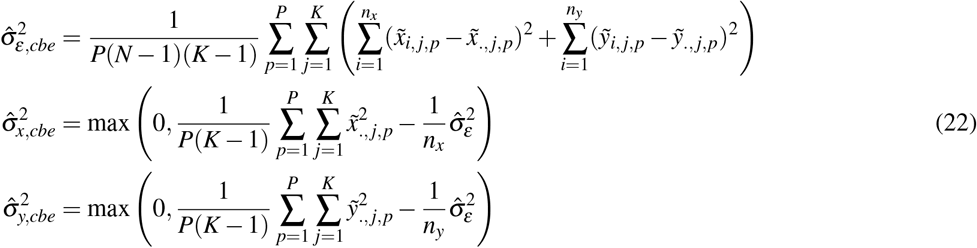

The estimator for the correlation then proceeds using Equations 13, 14, 15.

### 2.8 Group estimates

Given that the MLEs for correlations on individual datasets can become unstable, it is useful to obtain group estimates by assuming that some or all of the parameters are shared across subjects. For the group estimator explored in this paper, we assume that all parameters are the same across subjects. If the number of observations (*n*_*x*_ and *n*_*y*_) are also the same across subjects, this is equivalent to stacking all datasets along the voxel dimension, resulting in a single dataset with *P* = ∑ *P*^(*s*)^ voxels. In the more general case of unequal number of trials, we can maximize the sum of the log-likelihoods (Eq. 9) across subjects:

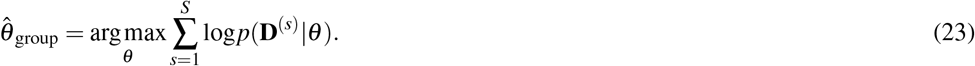

A corresponding cross-block group estimate can be obtained by computing the quantities in Eq. 12 and Eq. 13 across all voxels from all subjects, and then using these combined estimates to arrive at a correlation estimate (Eq. 15).

We also explored a maximum-likelihood estimator, for which we only assumed the correlation *ρ* to be the same across all subjects, but allowed each subject to have separate variance parameters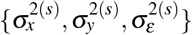.

### 2.9 Functional SNR

We defined the functional signal-to-noise (fSNR) of a multivariate pattern by the ratio of the variance of the true pattern and the variance of the measurement noise for the mean pattern estimate 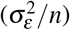 as

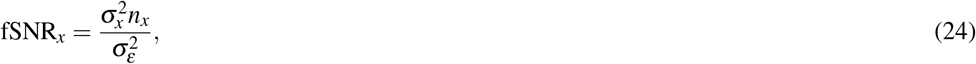

with an equivalent definition of fSNR_*y*_. In all simulations presented here, we used the same fSNR for both conditions. In additional analyses (see Appendix 6.5) we show that when the fSNR-levels of the two conditions differ by less than factor 7, the ML estimates behave relatively similar to simulations in which both fSNRs are equal and set to the geometric mean of the original fSNRs. As an empirical estimate of the overall fSNR, we therefore define

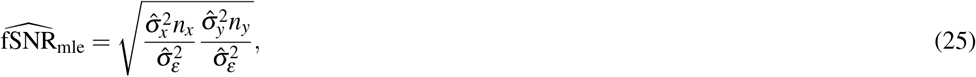

were all estimates are the maximum-likelihood estimates (mle). We can define a similar estimate using the cross-block estimates (cbe).

### 2.10 Simulation studies

We generated artificial datasets by drawing 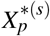 and 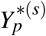 from independent normal distributions with mean 0. The signal variances were set to the same value for *X* and *Y*, and varied between exp(*−* 6) and exp(2). We generated *P* = 30 independent voxels for each individual dataset. Using Eq. 1, we then generated *n*_*x*_ = *n*_*y*_ = 6 independent observations for each condition. The measurement noise was drawn from a standard normal distribution, i.e. *σ*_*ε*_ = 1. Taken together, we therefore explored log(fSNR) values between *−* 4.20 and 3.79. For individual simulations (Fig. 1), we generated 5000 independent datasets with the true correlation set to *ρ* = 0.7. For group simulations, we simulated 5000 independent groups with *S* = 20 subjects each for each parameter settings. To assess the factors that influence the stability of the MLE, we repeated the simulations using 30, 150 or 750 voxels, and 4, 6, or 8 independent measures per condition.

### 2.11 Types on inference

Hypotheses about correlation coefficients fall into two categories. In the one-sample problem, we want to test whether the true correlation is larger or smaller than a specific value. In the simplest case, we want to test whether there is a positive linear relationship between two conditions, i.e. whether *ρ >* 0. Secondly, we want to determine whether the correlation is smaller than a high value, *ρ <* 0.99. A significant result in this one-sample problem would suggest that the two conditions engage partly separate processes in the region of interest, with the amount of separation specified by hypothesized correlation value. Third, we want to determine if the correlation is larger that than a specified value. A significant result would indicate that the overlap between two patterns has at least a specific size.

In the paired-sample problem, we are interested in comparing two correlations (i.e., *ρ*_1_ *> ρ*_2_) to determine whether they the correlation differs across two pairs of conditions or across two brain regions. Because fSNR can differ quite substantially between conditions and brain regions, we require stable estimates of the correlation coefficients. We will consider the one-sample and paired-sample situation in turn.

### 2.12 One-sample inference on correlations

We explored different procedures to test the hypothesis that the correlation is either smaller or larger than a specific value. First, we explored the use of the one-sided t-test and sign-test using individual correlation estimates. The test can be performed on the uncorrected 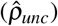 or maximum-likelihood estimates 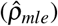.

To draw inferences on the less-biased group MLEs, we used a bootstrap (Efron et al., 1994) procedure to obtain confidence intervals around the estimate. In short, we drew 1000 bootstrap samples of size *S* = 20 with replacement from the original sample of the same size. We then obtained the group correlation estimate for each bootstrap sample.

To test whether the group correlation was smaller than a hypothesized value, we determined the proportion of bootstrap samples that were equal to or exceeded the hypothesized value, which provides a proxy for the *p*-value. To test whether the group correlation was larger than a specific value, we determined the proportion of bootstrap samples that were equal or smaller than the hypothesized value. This procedure is equivalent to constructing a (1*−* 2*p*) *×* 100% central confidence interval using the empirical quantiles of the bootstrap distribution and checking if the hypothesized value fell outside these confidence bounds.

To determine the validity of different tests, we simulated 5000 iterations of *S* = 20 subjects with true correlations of *ρ* = *{*0.7, 0.8, 0.9, 1.0*}*, and then tested against hypotheses of the same range of correlation values. All other simulation parameters were kept the same from the individual simulations. We measured Type-I error rates as the proportion of simulations for which the test was significant when tested against the true correlation. This proportion was determined separately for the two directions (smaller or larger) and for different *α*-value. We measured the power of the tests as the proportion of simulations for which the test was significant at a specific *α*-value when testing datasets against a hypothesized value that was larger or smaller than the true correlation.

### 2.13 Paired-sample inference on correlations

In paired inference, we seek to determine if the correlation between two activation patterns in one brain region is larger than in another brain region, or if the correlation between one pair of conditions is larger than between another pair of conditions. We assume that both brain regions or both pairs of conditions are measured in each subjects.

For inference based on individual estimates, we determined the correlation for each subject and region separately and then conducted a paired t-test across observations. For inference based on group estimates, we conducted a paired bootstrap. We generated 1000 bootstrap samples of size *S* = 20 from original sample of subjects with replacement, and then obtained group-correlation estimates for each region separately. For each bootstrap sample, we noted the difference in correlations 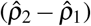. We then conducted a test by constructing the (1*− α*) *×*100% central confidence interval based on the quantiles of the bootstrap distribution. To test one-sided hypotheses *ρ*_1_ *< ρ*_2_ and *ρ*_1_ *> ρ*_2_, we determined whether 0 was below or above the interval, respectively.

To assess the validity of these procedures we simulated 5000 groups of *S* = 20. The log signal variance for one region (or pair of conditions) was varied from *−*3.5 to*−* 0.5, and for the other from *−*0.5 to *−*3.5, such that the average log(fSNR) across the two two regions was held constant at *−*2 + *log*(*n*). The first region was simulated with a correlation of *ρ*_1_ = 0.7, and the second region with a correlation *ρ*_2_ = 0.6, 0.7, or 0.8. We used 30 voxel for each region. To assess Type-I error rate and power, we then determined the number of simulations for which each test became significant at a specific *α*-level.

### 2.14 Empirical example study

As an empirical example, we utilized data from a published experiment investigating the planning and execution of simple and sequential finger movements (Ariani et al., 2022, 2024). On each trial, subjects prepared and subsequently executed either simple finger movements (digit 1,3, or 5), or sequential finger movements consisting of 6 presses (135315, 351531, or 513153). The preparation phase was 4-8s, followed in 60% of trials by a go-cue, and 40% of trials by a no-go cue. The activation pattern for preparation was estimated on no-go trials only, the execution-related activity only from go-trials (for details, see Ariani et al., 2022, 2024). Thus, the example was a study in the multi-pattern setting with 2 conditions (preparation and execution) and 3 items (the three movements). All procedures of this study were approved by the Ethics committee of Western University.

## 3 RESULTS

The results are structured as follows: We start by studying how uncorrected and maximum-likelihood estimates for individual datasets behave in the low signal-to-noise domain, and show that the maximum-likelihood estimate on a group of subjects is more stable than the average of the individual estimates. We then compare different inference techniques using these estimates, both for the one-sample problem, where we want to compare a single correlation against a specific value, and for the paired-sample problem, where we want to compare two correlations. We will discuss under what circumstances inferences are valid, and when they are not. Finally, we apply the method to real datasets to show the utility for answering neuroscientific questions.

### 3.1 Individual estimates

To study the behavior of different correlation estimates, we simulated data with a known correlation of *ρ* = 0.7 for a range of functional signal-to-noise ratios (fSNR, see methods) that are typical for unsmoothed individual-voxel fMRI data.

As the fSNR decreased, the uncorrected estimate (Fig. 2a, dashed line) systematically underestimated the true correlation, as expected. In contrast, the MLE corrected for the bias effectively for log(fSNR) values above *−* 0.5. For lower fSNR, however, the MLE also started to show a bias. Additionally, the median of the MLEs first became positively biased, before converging towards zero for very low fSNRs. To understand this complex behavior, we need to consider that the MLE is constrained by multiple boundaries (Fig. 3a). The correlation estimate is bounded between 1 and *−* 1 and the estimated true pattern variances are bounded by zero - meaning that the overall estimated fSNR cannot be negative. For lower fSNRs, the correlation estimates increasingly approached the boundaries at | *ρ*| = 1. The number of cases approaching the upper boundary considerably outweighed the number of cases approaching the lower boundary, causing the median estimate to be higher than the true value. For very low fSNRs, the estimated pattern variances often approached zero 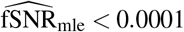. The proportion of these cases increased with decreasing fSNR and reached 40% for pure noise data (see Fig. 2b). Finally, for pure noise data, the correlation estimate was equally likely to be 1 or *−*1, such that both in mean and median of the MLE approached zero.

**Figure 2.**
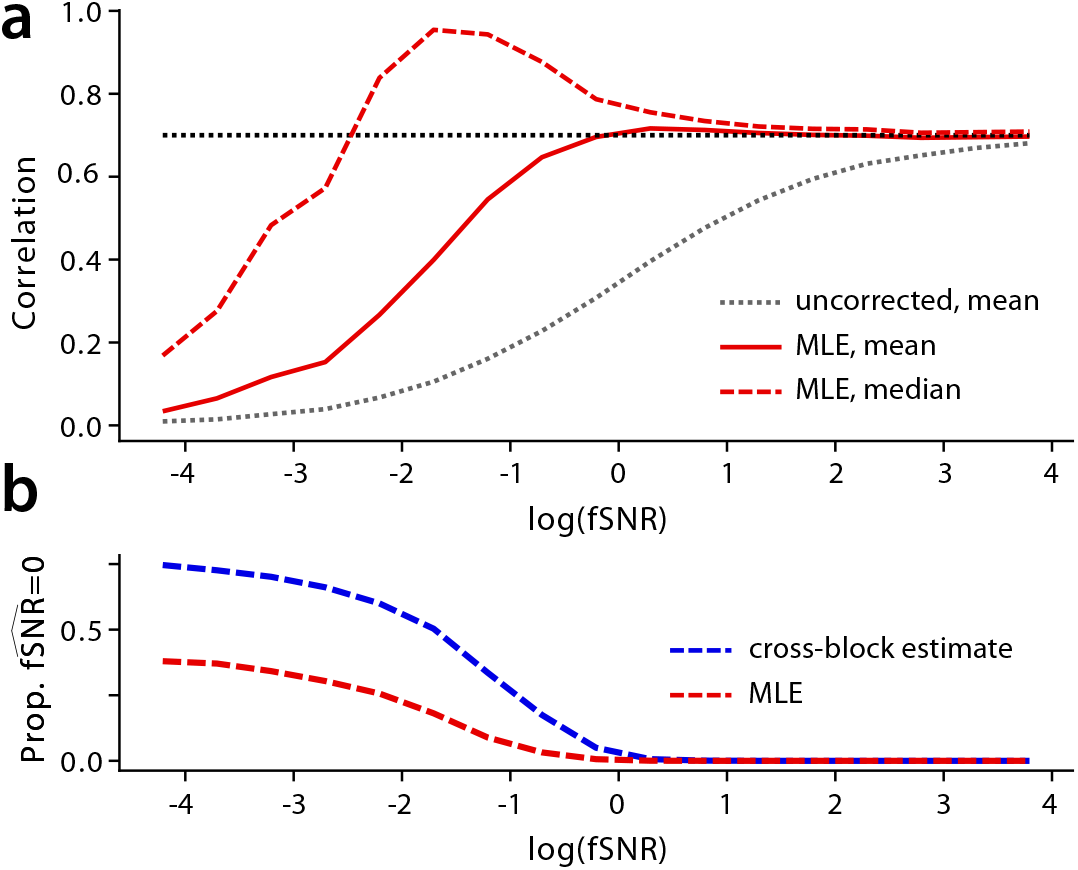
Correlation estimates from individual datasets, simulated with a true correlation of *ρ* = 0.7. **(a)** Mean (solid line) and median (dashed line) of maximum-likelihood correlation estimates as function of log(fSNR). The uncorrected estimate is shown in the gray dashed line. **(b)** Proportions of cases in which the fSNR was estimated to be zero for the cross-block (blue) and the maximum-likelihood (red) estimate.

**Figure 3.**
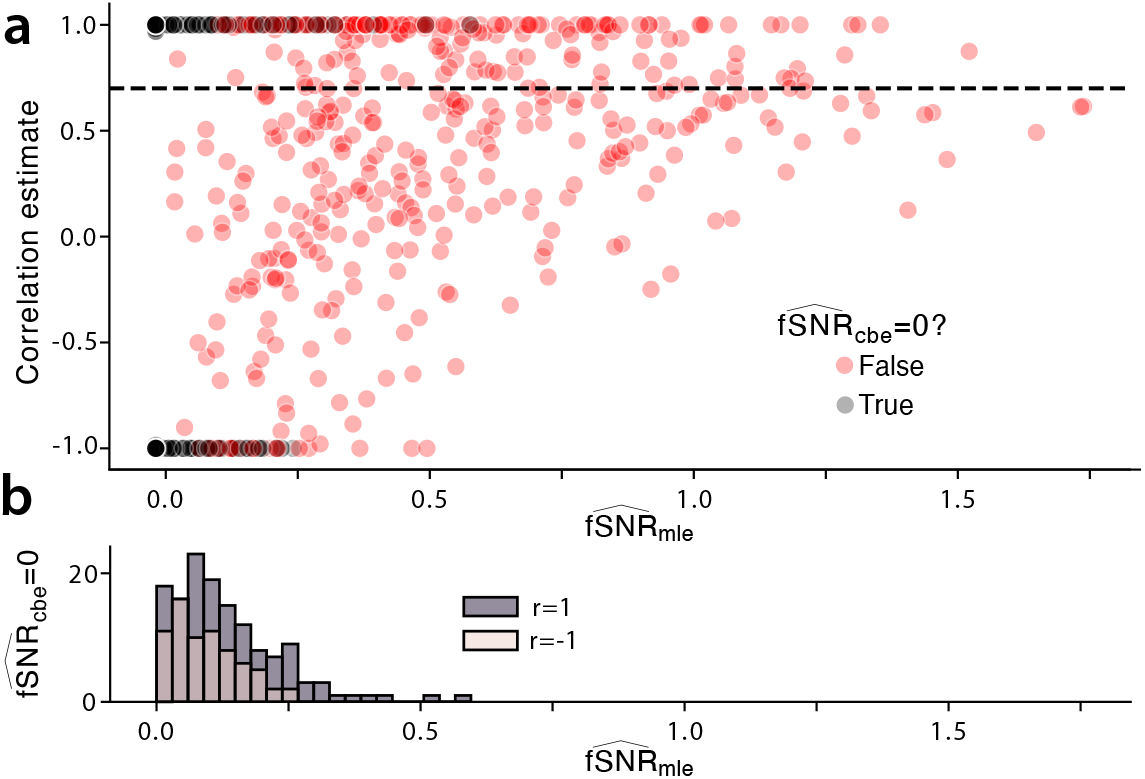
Behavior of the maximum-likelihood estimate for low fSNRs. **(a)** Maximum likelihood estimate of correlation 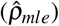 plotted against estimated 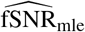. The datasets were simulated with a true correlation of *ρ* = 0.7 and log(fSNR) between *−*3.2 and *−*0.2. Black dots indicate cases for which 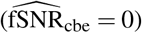. **(b)** Number of datasets out of 800 simulations for which 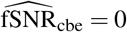, but 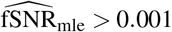.

For low fSNRs, the MLE and the cross-block estimates also behaved differently. While the correlation estimates for the two approaches were very close, the variance estimates could differ substantially. Specifically, when the correlation estimate was close to a bound, i.e. |*ρ*| = 1, the 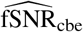 often became zero, even though the corresponding MLE was clearly positive 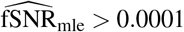. This occurred quite often (Fig. 2 a, b) such that with decreasing true fSNR the number of simulations with 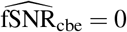 reached 75% (Fig. 2b).

In summary, the cross-block and maximum-likelihood approaches resulted in very similar correlation estimates. If we are using all correlation estimates, then the two approaches can be used interchangeably. However, if we wish to exclude estimates for which the estimated fSNR is zero, then cross-block estimate will lead to roughly twice as many exclusions as the maximum-likelihood approach. We will consider the question of exclusion of estimates for inference, showing that it is generally advantageous to retain all estimates.

### 3.2 Bias and stability of estimates

To understand at what point the maximum-likelihood estimate starts to show a considerable bias, we repeated the simulation, this time also varying the number of independent measures (*n*_*x*_ = *n*_*y*_), and the number of independent voxels (*P*).

For the uncorrected correlation estimate, the bias depended only on the fSNR (a function of the signal variance, noise variance, and number of measures, see methods), but was independent of the number of voxels (Fig. 4a).

**Figure 4.**
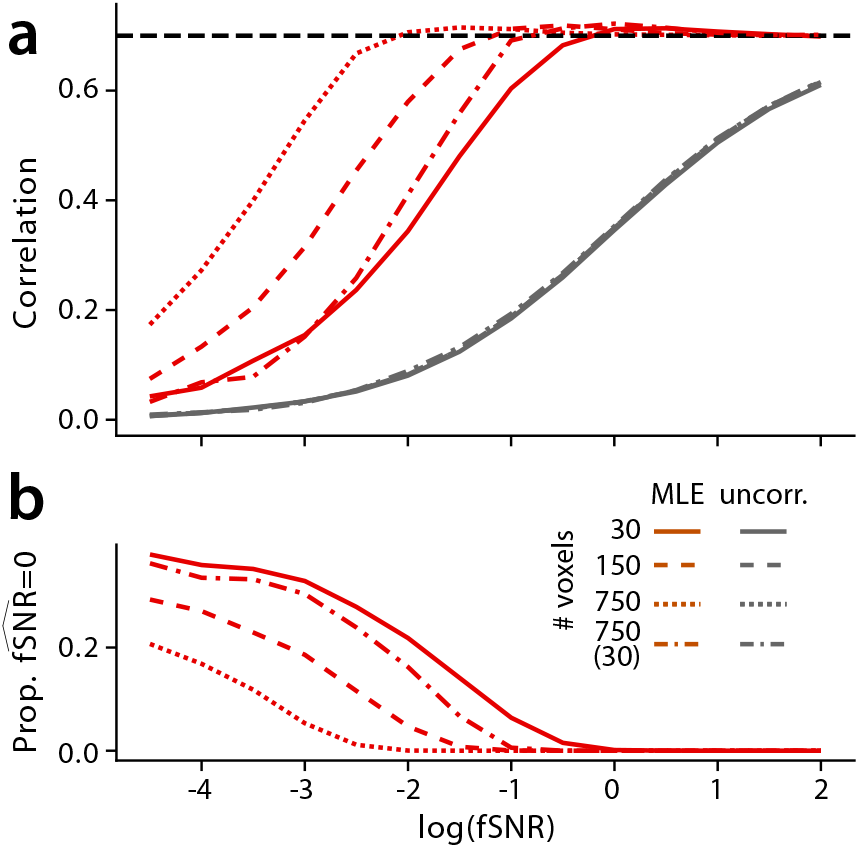
Correlation estimates from individual datasets using different number of voxels and fSNRs. **(a)** Mean of uncorrected (gray) and full maximum-likelihood estimate (red) as a function of number of independent voxels and log(fSNR). The dash-dotted line shows a simulation with 750 voxels with spatially correlated noise and effective number of voxels of *P*_eff_ = 30. **(b)** Proportion of datasets with 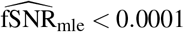.

In contrast, the behavior of the MLE depended both on fSNR and the number of independent voxels. As the number of independent voxels increased from 30 to 150 to 750, the point at which the estimate shows substantial biases moved to lower fSNR levels. The bias arises from the fact that the corrected variance estimates (Eq. 12) have high variance - and this variance decreases when more voxels are available.

These insights have two important practical consequences when trying to obtain stable correlation estimates. First, given a fixed amount of data, we cannot improve our estimate by averaging the patterns across observation. While each pattern estimate becomes less noisy, the overall fSNR remains unchanged. That is, in general we recommend to use all independent measures for *x* and *y* that are available. Secondly, we can improve the stability of correlation estimates by using more voxels in the estimation. While using a larger region engenders a loss of spatial specificity of our conclusion, it is generally best to use the largest possible spatial scale for which the inference is still meaningful.

### 3.3 Spatial dependence of voxels

The stability of the MLE depends on the number of independent voxels. In fMRI data, however, spatially neighboring voxels show strong noise correlations that arise from both in intrinsic spatial smoothness of the noise processes, as well as interpolation during data pre-processing (Arcaro et al., 2015; de Zwart et al., 2008). It can be theoretically shown (Diedrichsen et al., 2021) that the variances of the signal variance and covariance estimates (Eq. 12, 13) depend on the *P× P* covariance matrix of the measurement noise (Σ_*P*_). Specifically the variance of the cross-block estimate of the variances scales with

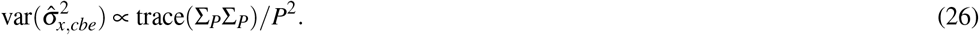

For independent voxel 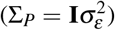 this quantity simplifies to 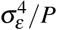. For dependent voxels it can be expressed as 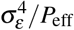, with the effective number of voxels (*P*_eff_) being

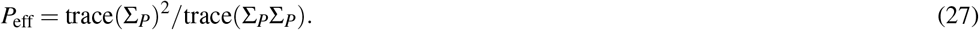

To test whether this scaling of the variance generalizes to the distribution of correlation estimates, we repeated our simulation with *P* = 750 voxels, but a correlation structure such that *P*_eff_ = 30. The bias of this estimate as a function of fSNR was indeed similar to that observed for 30 independent voxels (dash-dotted line in Fig. 4), even if the behavior was not identical.

In practice, we can bring the measurements closer to the assumption of identical and independent noise by either univariate (dividing each voxel by a estimated noise standard deviation) or multivariate pre-whitening (post-multiplying the pattern estimates with and estimate of 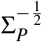. Both approaches have been shown to reduce the variance of variance and covariance estimators (Walther et al., 2016), and will therefore increase the stability of the correlation estimates. It should be noted, however, that even multivariate pre-whitening never fully makes the voxels independent, as we do not have access to the true noise covariance matrix.

As a consequence, it is difficult to determine what combination of fSNR and number of voxels is sufficient to obtain stable estimates of the correlation of two patterns. We therefore suggest a simple criterion based on the distribution of the MLE estimates themselves (see discussion).

### 3.4 Group estimates

When estimating correlations for a group of datasets (subjects), we can average the MLEs obtained on each individual subject. Using individual estimates has the potential advantage that we can draw inferences at the group level using standard statistical tests (i.e., t-test). As we have seen, however, individual estimates are unstable and biased for low fSNR levels. The group estimates obtained by averaging valid independent individual estimates (Fig. 5) therefore shows the same strong bias for uncorrected estimates (gray dotted line), and for lower fSNR levels also for the MLE (red dotted line).

**Figure 5.**
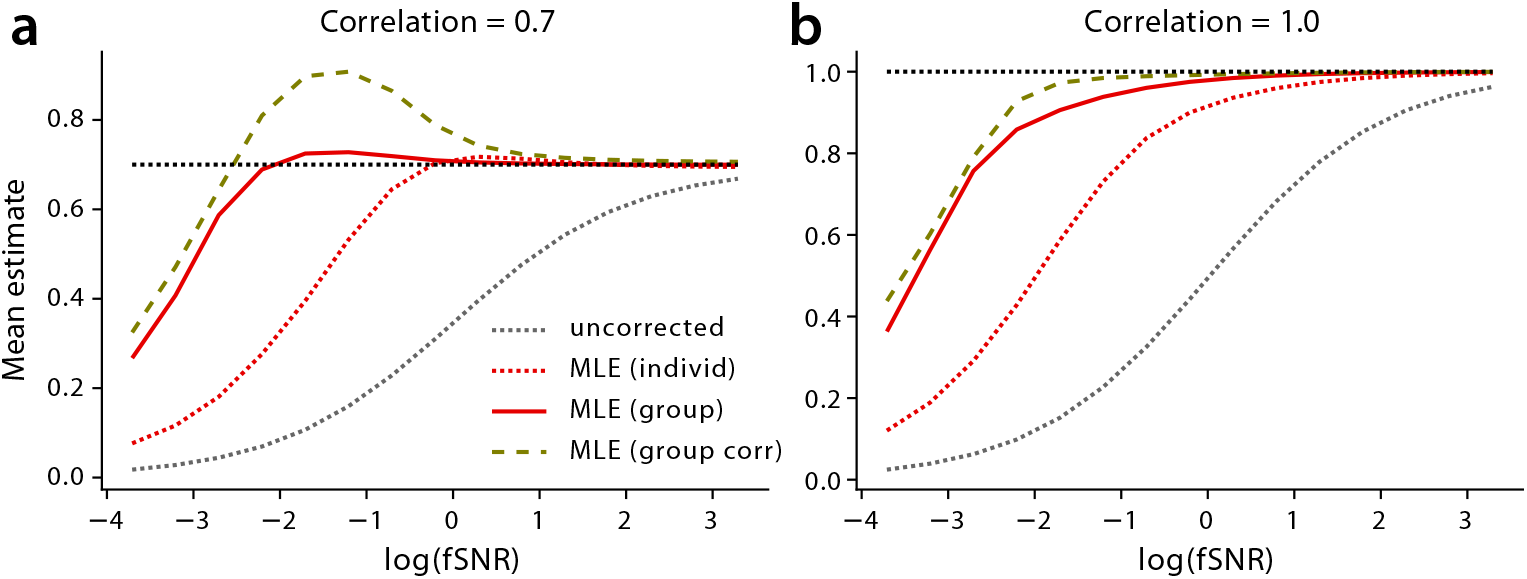
Correlation estimates for a group of *N* = 20 datasets. Mean group estimate, derived from averaging individual estimates (red and gray dotted lines), or by fitting a group models with shared parameters to all datasets (MLE, group, solid line). The dashed line (MLE, group corr.) shows a model for which the correlation parameter is shared, but variance parameters are individual. The x-axis shows the log(fSNR). **(a)** True correlation is *ρ* = 0.7 (left), or **(b)** *ρ* = 1.0.

To derive a more stable group estimate, we may want to assume that the parameters are the same across all individuals, which allows us to combine all data when maximizing the likelihood (see methods). This is effectively the same as concatenating the voxels patterns of all individuals along the voxel dimension, which increases the number of voxels. Similarly to using more voxels (Fig. 4), this increases the stability of the correlation estimates down to very low fSNR levels (red solid line).

We also explored a maximum-likelihood estimator, for which the correlation *ρ* parameter was the same across all individuals, but where the variance parameters 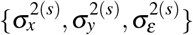 were specific to each individual. While this estimator (Fig. 5, dashed line) performed well when the true correlation was *ρ* = 1.0 (right), it exhibited a strong positive bias for intermediate fSNR levels for smaller correlations (*ρ* = 0.7, left). For inference, we therefore only considered the group MLE with all parameters shared across individuals.

### 3.5 One-sample inference

In the one-sample problem, we test whether the correlation is smaller or larger than a specific value. In the simplest case, we want to test whether the correlation is larger or smaller than zero. In more complicated cases, we want to test if the true correlation is smaller than a specified value (for example *ρ <* 0.99), or larger than a specified value (for example *ρ >* 0.8). These tests can help us establish the exact degree of overlap between two patterns of activation.

We considered a range of methods to test these hypotheses. First, we can use individual uncorrected (Pearson) correlation estimates and conduct a one-sample t-test against the hypothesized value. Alternatively, we can use the same approach using the maximum-likelihood estimates. When testing against the simple null hypothesis that *ρ* = 0, both approaches controlled well for the Type-I error rate (Fig. 6a). When we tested how well these approaches can detect a true correlation of size *ρ* = 0.2, we found that both estimates had similar power (Fig. 6e).

**Figure 6.**
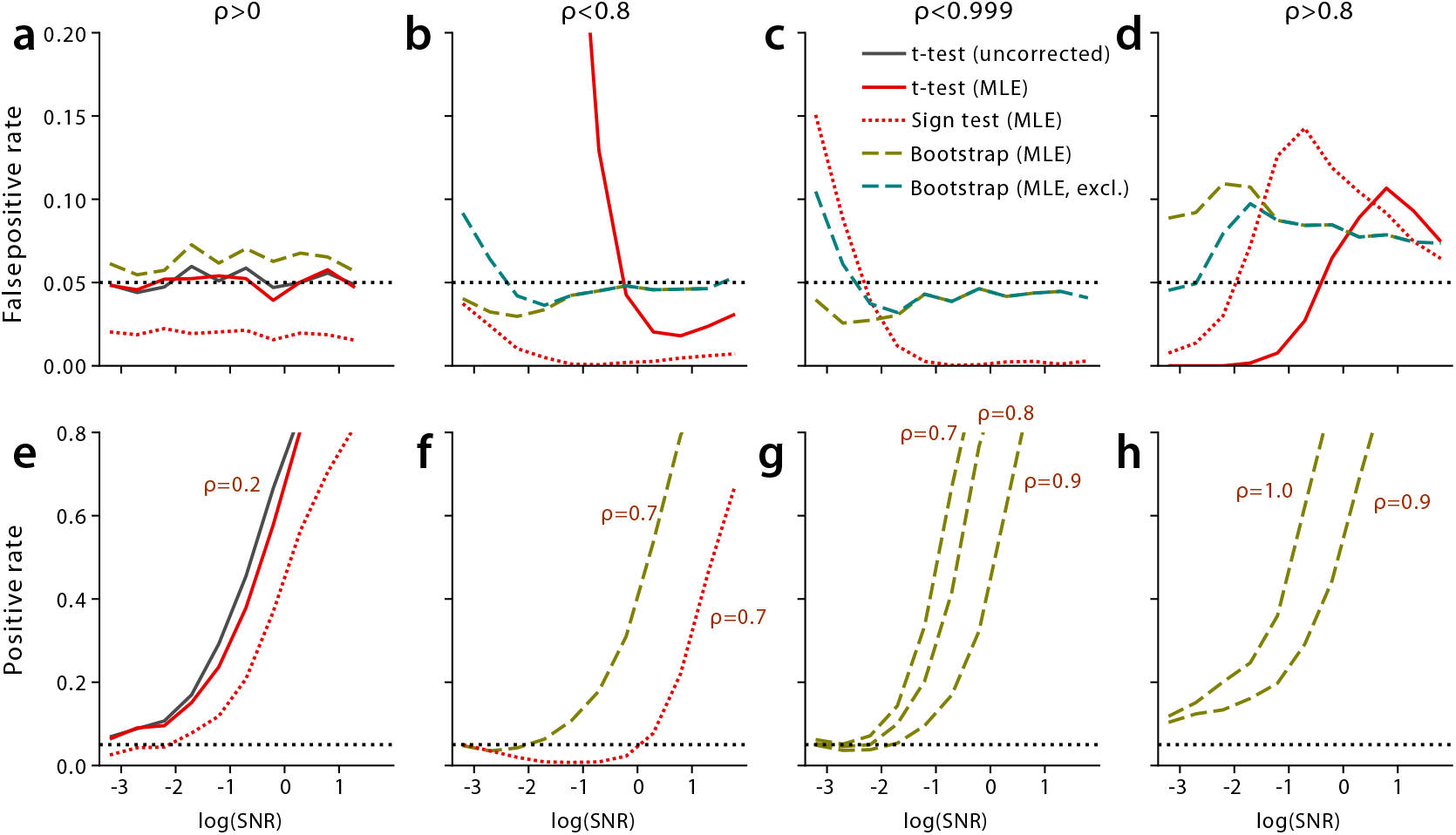
Positive rate for the one-sample problem. The correlation of samples of *S* = 20 subjects is tested (from left to right) to be larger than zero, smaller than 0.8, smaller than 0.999, or larger than 0.8. **(a-d)** False positive rate for different methods for *α* = 0.05 (dashed line) when the data is simulated under the Null-hypothesis. T-test and sign-tests are conducted on individual MLEs. The bootstrap uses a group estimate (full MLE or cross-block estimate). **(e-h)** Power (true positive rate) when the correlation of the simulation is set to the value indicated next to the line.

However, when testing whether the true correlation is smaller than a set value, the use of individual estimates failed. The Type-I error rate for the uncorrected estimates was close to 100% when testing whether the correlation was smaller than a specific value, and 0% when testing whether the correlation were larger than a specific value. The use of the less-biased MLE did not fix this problem. For testing *ρ <* 0.8, the Type-I error rate exceeded the set *α*-level of 0.05 for low fSNRs (Fig. 6b). For the hypothesis of *ρ <* 0.999, the t-test using the MLE was virtually always significant, with Type-I error rates exceeding 80% (not shown). This is due to the fact that the estimates are bounded at 1.0 and slightly downward biased (Fig. 5). For testing whether *ρ >* 0.8, the Type-I error exceeded the set value for large fSNR values, before going to zero. That is, in all these cases, the bias of the individual MLEs prevents valid inference using a t-test on individual estimates.

Alternatively, we could consider using a sign-test, counting the number of individual estimates that exceeded a certain value, and assessing the probability of this (or a more extreme) outcome on a binomial distribution. For testing hypotheses *ρ >* 0 and *ρ <* 0.8, this method did control the Type-I error rate (Fig. 6a,b, dotted line). This, however, came at the expense of statistical power (Fig. 6e,f). When testing *ρ >* 0.8, the Type-I error rate exceeded the set value. Therefore, the bias of the median of individual maximum-likelihood estimates also renders the sign-test virtually useless.

As an alternative, we therefore turned to the more stable group MLE, and used a subject-wise bootstrap to obtain confidence bounds for our estimates that takes into account the subject-by-subject variability of the correlation coefficient. In short, we iteratively resampled *S* subjects with replacement from the original sample of *S* subjects, each time obtaining a (fixed effect) group MLE of the correlation (for details see methods). For testing the hypotheses *ρ <* 0.8 and *ρ <* 1.0 this method effectively controlled the Type-I error rate at the required level, even for very low fSNRs (Fig. 6a, red dashed line). Even though the estimates are bounded at *ρ <* 1, we can test the hypothesis *ρ <* 1.0: As long as at least a proportion of *α/*2 bootstrap samples fall on that boundary, the (1*− α*)100% central confidence interval will include 1.0, rendering the test non-significant.

For the bootstrap, we also tested the effect of excluding correlation estimates for which the cross-block fSNR estimate is 0. For low fSNR values, this procedure led to a Type-I error rate that exceeded the desired level for testing whether the correlation is smaller than a set value (Fig. 6b,c). For pure noise data (fSNR = 0), the Type-I error for the bootstrap with exclusion led to a Type I error rate of 34.4% (not shown). In contrast, the bootstrap using all MLEs, controlled the false positive rate better, although not perfectly. For example, on pure noise data the Type-I error rates for *ρ <* 0.999 exceeded the 5% level in 8% of the simulations. Finally, the bootstrap method showed good power for testing these two hypothesis (Fig. 6f,g). It started to detect the alternative hypothesis at low fSNR, with power rapidly increasing for higher fSNRs.

Only when testing whether a correlation is larger than a specific value, for example *r >* 0 (Fig. 6a) or *ρ >* 0.8 (Fig. 6d), did the bootstrap not behave as desired. The Type-I error rate exceeded the required level, reaching Type-I error rates of nearly 0.1 for *α* = 0.05. Nonetheless, compared to the other methods, this behavior was relatively stable across fSNR levels. In conclusion, the subject-wise bootstrap results in confidence intervals for which the upper bound is usually adequate, but the lower bound tends to be too high.

### 3.6 Paired-sample inference

Finally, we considered the problem of testing whether a pattern correlation differs between two regions, or between two sets of conditions within the same region. The problem when testing this hypothesis is that different brain regions (or different conditions) can have very different fSNRs.

To test this scenario, we simulated data for two regions (or two different sets of conditions) for *S* = 20 subjects. We then varied the log(fSNR) for the two regions in opposite directions, such that the average log(fSNR) was always *−* 0.3. The correlation of the first region was always *ρ* = 0.7. We first assessed how different methods performed for testing the hypothesis *ρ*_1_ *> ρ*_2_ in the case when the data was generated under the Null-hypothesis *ρ*_1_ = *ρ*_2_ = 0.7.

As expected, a paired t-test on uncorrected Pearson correlation (Fig. 7a, dark gray line), was very strongly biased by the fSNR difference. The test for *ρ*_1_ *> ρ*_2_ was nearly always significant when fSNR_1_ *>* fSNR_2_, and nearly never significant when fSNR_2_ *>* fSNR_1_. A paired t-test on individual maximum-likelihood estimates (red line) showed the same bias, albeit the Type-I error rates were less severe. Nonetheless, unless the SNR values across the two sets are nearly identical, individual correlation estimates cannot easily be compared.

**Figure 7.**
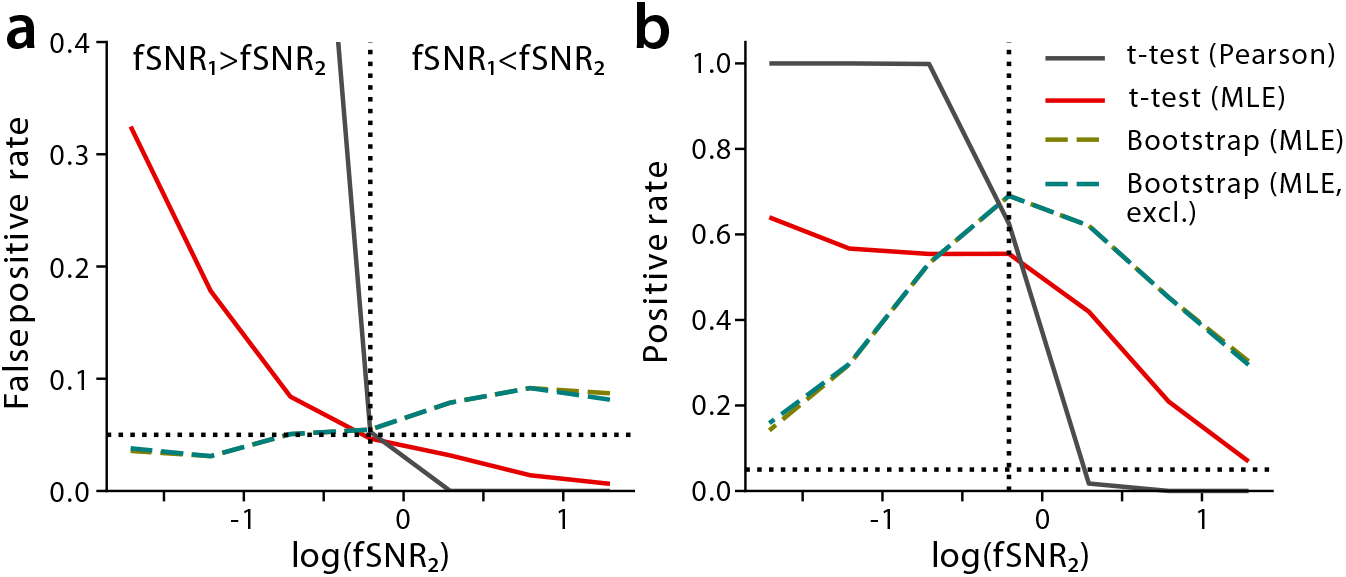
Positive rate for a paired-sample test of the hypothesis *ρ*_1_ *> ρ*_2_ with unequal fSNR. **(a)** False positive rate when the data was simulated under the Null-hypothesis *ρ*_1_ = *ρ*_2_ = 0.7. **(b)** Power (true positive rate) when *ρ*_1_ = 0.7 and *ρ*_2_ = 0.45. Horizontal dashed line indicated the desired *α* = 0.05 level, vertical dashed line the point where fSNR_1_ = fSNR_2_. For the hypothesis *ρ*_1_ *< ρ*_2_ we obtain symmetrical results (not shown).

In contrast, the bootstrap on the group estimates (Fig. 7, dashed lines) was more stable across a range of fSNR values. For the critical domain that fSNR_1_ *>* fSNR_2_, the Type-I error was well controlled. In this domain the test also still had reasonable power to detect that *ρ*_1_ *> ρ*_2_ when *ρ*_2_ was set to 0.45 (Fig. 7b).

However, given that the lower bound of the bootstrap interval tends to be too high (see one-sample inference), the test was not perfect: The test declared the correlation in the set with lower fSNR to be bigger more often than the set *α*-level of 0.05. Therefore, in this domain we need to exercise some caution in interpreting significant results.

### 3.7 Multi-pattern setting

The methods introduced here also apply to the setting in which have multiple items per conditions and want to test whether these items are “represented” in a similar way across conditions. In essence, we want to test whether the differences between the different items are parallel (*ρ* = 1) or orthogonal (*ρ* = 0) across conditions (Fig. 1c). We can test this idea by removing the mean pattern for each condition, and assessing the correlation between the vectors connecting each item with the condition mean across the two conditions.

To deal with this problem, we require some modification that accounts for the removal of the mean condition pattern (see method). Given these modifications, the methods and results for the single-pattern setting fully generalize to the multi-pattern setting. As can be seen in a simulation with 4 items per condition (Fig. 8) the MLE again effectively corrects for the negative bias until the fSNR becomes too low. In the multi-pattern setting, the stability of the estimate is determined by the fSNR times the number of voxels, repetitions, and items.

**Figure 8.**
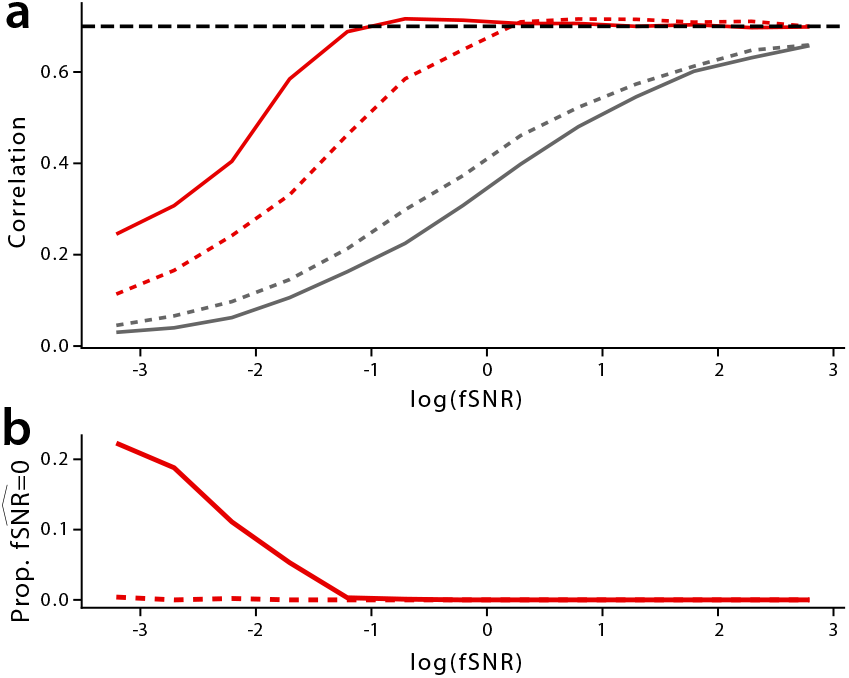
Individual estimation using multiple items per condition. **(a)** Mean uncorrected (gray) and maximum-likelihood (red) correlation estimate for a simulation with 4 items per condition. Either all voxels (solid line) or only voxels with positive reliabilities across partitions (dashed line) were use. **(b)** Proportion of 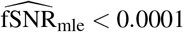 without (solid line) and with (dashed line) voxel selection.

### 3.8 Effects of voxel selection

Overall, the fSNR in multivariate fMRI studies is often quite low. This issue arises especially in the multi-pattern setting, in which the differences between different items within a condition can be quite subtle. In such cases, the researcher may be tempted to estimate correlations only on informative voxels within a region. Unfortunately, this approach induces downward bias of the MLEs. To demonstrate this, we simulated voxel selection by only including voxels that had positive signal variance estimates for *X* and *Y*. In this case, the signal variance was defined as the variance of the true activation values for each voxel across items (see method). While such selection slightly improved the quality of the uncorrected correlation estimate, it severely biased the MLE (Fig. 8, dashed lines). This is because the signal-variance estimate in each condition is uncorrelated with the covariance estimate across conditions. By truncating voxels with low signal variance estimates, we overestimate the signal variance, while leaving the average covariance estimate unchanged, causing us in turn to underestimate the correlation. When testing the correlation against a fixed value (see above) this procedure therefore would lead to the erroneous conclusion that the two activation patterns (or multi-item representations) are more distinct than they actually are.

### 3.9 Real dataset examples

To show the utility of the proposed methods for real fMRI data, we utilized a multi-variate example from an experiment investigating the planning and execution of a single or sequential finger movement (Ariani et al., 2022, 2024). In short, subjects prepared and subsequently executed either 6 repeating single-finger movements (digit 1,3, or 5), or sequential finger movements consisting of 6 presses of different fingers (see methods). We found that there was information about the upcoming movement (both single and sequential) in the hand area of primary motor (M1) and somatosensory (S1) cortex. We then wanted to determine to what degree the pattern that indicates that a specific movement is planned correlates with the pattern observed during the execution of the same movement. For the sequential movements we hypothesized that planning would activate the first digit and thus would be most similar to execution pattern of the first finger in the sequence (Yokoi et al., 2018). Thus, we had here a multi-item design with two conditions (planning and execution) and three items each (finger movement).

We first analyzed the functional SNR in each region and condition after removing the mean pattern for execution or planning. While our simulations have always assumed that the fSNR is the same across the two conditions, this was not the case in the empirical example: the fSNR for execution was higher than during planning, with a log(fSNR) difference of 0.76. Given that the fSNR ration was less than 7 (see Appendix 6.5), the overall effective log(fSNR) was -1.59 for single finger planning and -1.54 for sequence planning. At these signal-to-noise levels, the uncorrected correlation estimate (Fig 9b) was strongly biased towards zero. Using these estimates, we could only establish that the correlations were larger than zero for each region and condition *t*_22_ *>* 3.057, *p <* 0.006.

**Figure 9.**
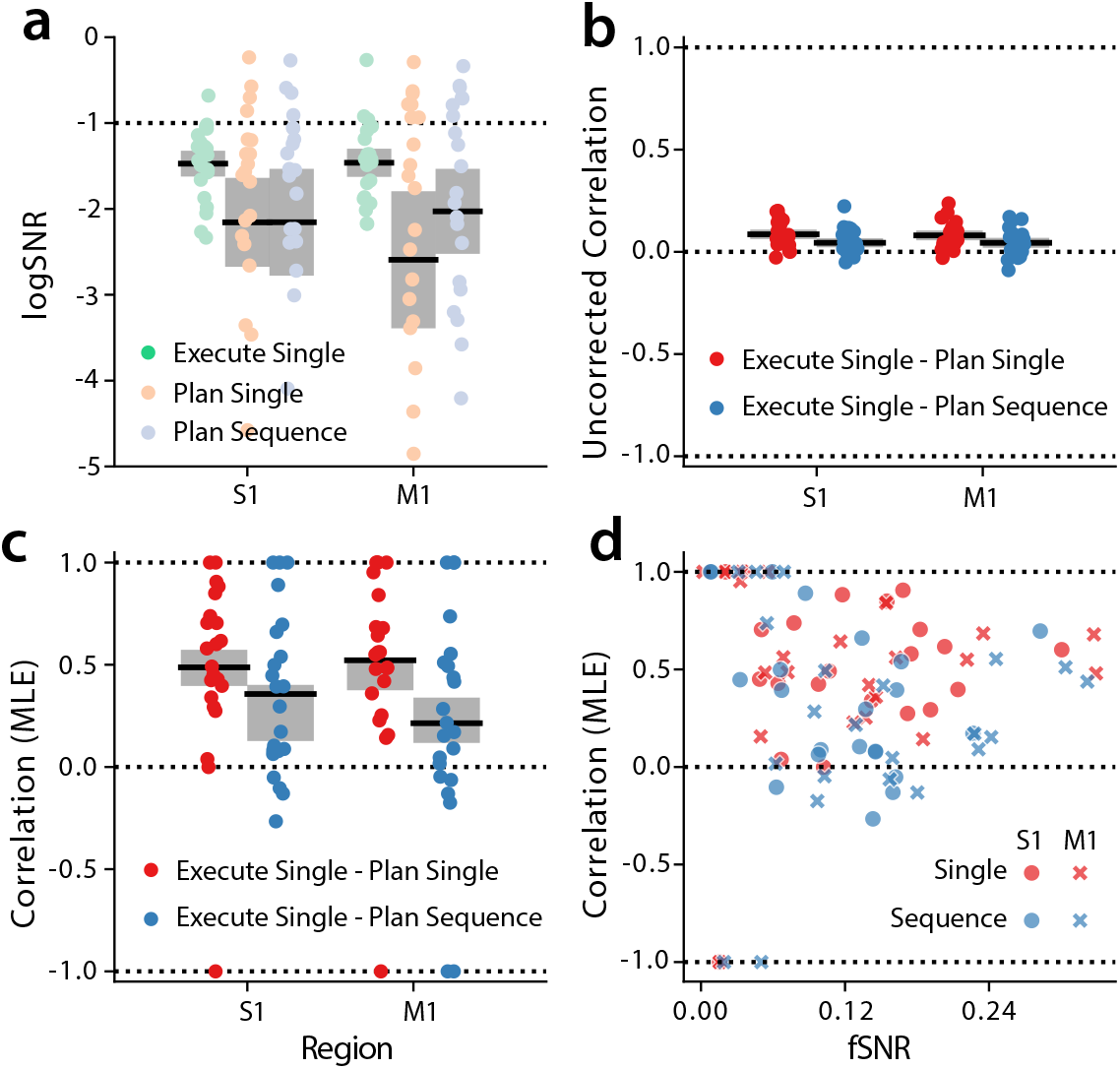
Application example to a study of planning and execution of individuated finger movements. **(a)** log(fSNR) for the execution of individual finger movements (green), planning of single finger movements (red), and planning of finger sequences (blue) in primary motor (M1) and primary somatosensory (S1) cortex. Each dot indicated an individual subject, solid line the mean, and gray box 90% confidence interval. **(b)** Uncorrected correlation coefficient between the activation pattern for planning and executing single finger movements (red), or planning sequential finger movements and executing a single finger movement with the same first finger as the sequence (blue). **(c)** Individual MLEs of correlations (dots) and 90% confidence interval based on group estimates and subject-wise bootstrap. **(d)** Individual MLEs of correlations as a function of the estimated fSNR in each subject.

To test how large the true correspondence between planning and execution activation patterns was, we derived the MLE (Fig 9c). Given that the ROI consisted of 526 (M1) and 1083 (S1) voxels, 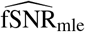 exceeded 0.0001 for every single subject, region, and condition. Nonetheless, as the fSNR estimate for individual subjects decreased, the variance of the correlation estimate increased with estimates increasingly falling onto the boundaries (Fig 9d). In comparision, the 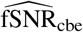 estimate was zero in 15.9 % of the subjects in M1 and 11.4% for S1.

We then determined the group estimate of the correlation and the corresponding 90% confidence interval using the subject-wise bootstrap. Based on the bootstrap distribution, we can conclude with *α* = 0.05 for a one-sided test that the true correlation was below *ρ* = 0.6, even for simple movements. This shows that, although there was considerable overlap between the patterns associated with planning and executing the same movements, the two did not correspond perfectly, but had a unique component in each condition.

The results also suggested that the correlation for single finger planning was higher than for sequence planning. In this case, the fSNR for sequences was higher than for single fingers, such that we are testing *ρ*_1_ *> ρ*_2_ in a domain in which fSNR_1_ *<* fSNR_2_. In this case the t-test on individual Pearson correlations is very conservative. Nonetheless, the difference in fSNR was small enough, such that even the t-test on uncorrected correlations was significant (S1: *t*_21_ = *−* 2.629, *p* = 0.0078, M1: *t*_21_ =*−* 2.064, *p* = 0.0258). When we corrected for the difference in fSNR by using the group MLE and bootstrap, we also found that the difference was highly reliable (S1: *p <* 0.0002, M1: *p <* 0.0002). Using this convergent evidence, we can therefore be relatively confident that planning the single finger movements overlaps more with the execution of a single movement than with planning of a sequence beginning with the finger - suggesting that the preparatory state for a sequence contains slightly more information than the first finger.

## 4 DISCUSSION

In this paper, we present a general form of the MLE of the true correlation between two variables measured with noise (Azen and Reed, 1973). As our main application of interest is to determine the correlation between two brain activation states from noisy fMRI data, we were especially interested in the behavior of these estimates and the associated inferential procedures for data with very low signal-to-noise ratios. We show that the MLE corrects for the strong bias in the Pearson correlation efficiently if a large number of voxels are available and the signal-to-noise ratio is still sufficient. For lower number of voxels or lower fSNR, the sampling distribution of the MLE concentrates near the boundaries of 1 and -1. We also find that the cross-block correlation estimator (Beaton et al., 1979; Saccenti et al., 2020; Liu et al., 1978) is nearly identical to the MLE. If cross-block estimated falls exactly on the bounds, then the MLE approaches numerically that that bound. In contrast, the estimated variances differ between the cross-block and maximum-likelihood methods when the estimate for the correlation falls on one of the two boundaries. Thus, for valid estimation of the fSNR, the maximum-likelihood approach is necessary. Another advantage of the MLE is that the model formulation can be easily adapted to novel situations, such as addition of fixed effects, different noise-variances across conditions, or multiple items per condition.

To test the hypothesis that the correlation is smaller than a specific value, we show we can draw valid inferences using a subject-wise bootstrap on the group MLE estimate. In contrast, the same method leads to slightly inflated Type-I errors when testing whether the correlation is larger than a specific value, indicating that the lower bound of the confidence interval is slightly too high.

Generally, we find, however, that the bootstrap of the group estimate degrades relatively gracefully: When the fSNR approaches 0 (pure noise), the bootstrap confidence interval includes *−*1 and 1 in most cases (if rounding six decimal places). Nonetheless, the rate of Type-I errors for testing the hypothesis *ρ <* 1 is typically slightly higher than the desired significance level, such that it is likely advisable to not over interpret results from regions where the fSNR is estimated to be very low. This is especially important when we test these hypotheses in many locations across the entire brain in a search-light mapping approach (Kriegeskorte et al., 2006), as it will result in many false positives and requires a strict correction for multiple testing (Friston et al., 1994).

### 4.1 Best practices when testing correlations in fMRI data

In summary, our results suggest the following best practices when testing hypotheses about correlation of brain activation pattern:

- As a diagnostic, we recommend plotting the individual maximum-likelihood correlation estimates against the estimated fSNRs (as in Fig. 9d). For the latter, we recommend using the geometric mean of the estimated fSNRs for the two conditions, or if the two differ by more than 7-fold (see Appendix 4), the lower of the two. If more than half of the fSNR estimates are close to the parameter bound of zero, or the correlation estimates spread evenly across the two bounds of *ρ* = *−*1 and *ρ* = 1, then the overall fSNR is likely too low to draw any reasonable inferences.
- When testing the hypothesis *ρ >* 0 or *ρ <* 0, simply conduct a one-sample t-test of the individual estimates against zero. This can be done on the uncorrected Pearson correlations (or cosine similarities) or using the MLEs.
- When testing the hypothesis *ρ < x* (with *x* being a positive number), we recommend estimating the group MLE and conducting a subject-wise bootstrap to obtain confidence bounds on the estimate. For a valid bootstrap, the sample size should be at least *S* = 20. The p-value can then be determined by the proportion of bootstrap estimates that exceed or are equal to *x*.
- When testing the hypothesis *ρ > x* (with *x* being a positive number), the bootstrap procedure tends to have a Type I error rate that exceeds the desired value by a little less than factor 2. We therefore recommend correcting for this by choosing a twice as stringent statistical threshold.
- When testing the hypothesis *ρ*_*>*_*ρ*_2_ across two regions or two different sets of conditions, first establish the relevant fSNR values for the two cases. This is either the geometric mean of the two fSNRs, or the lower of the two, if they differ by more than factor 7 (see Appendix 4).
  – If *SNR*_1_ *> SNR*_2_ (the bias in individual correlation estimates favors the alternative hypothesis), use a paired bootstrap on group MLEs to draw inferences.
  If *SNR*_1_ *< SNR*_2_ (the bias in individual correlation estimates favors the Null hypothesis), you can use a paired bootstrap with a twice as stringent significance threshold. Convergent evidence can be found by using a paired t-test using the individual maximum-likelihood estimates, which is conservative in this setting.

If the fSNR of the data is too low to draw valid inferences, we recommend the following approaches:

- Consider increasing the size of the region. This will have some limits, as we are often interested in brain regions of a specific size. As a general rule, however, it is advisable to choose the coarsest level that still allows answering the scientific question at hand.
- Reduce spatial noise correlation by multivariate pre-whitening the data (Walther et al., 2016). Appropriate regularization to the estimated noise covariance matrix 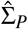_*P*_ has to be applied. This step often improves the stability of the correlation estimates by increasing the effective number of voxels (Eq. 27).
- If both approaches fail, the solution may simply lie in getting more and better data. Acquiring data with higher spatial resolutions can help, as the number of available voxels increases. However, this direction has limits, in that it also decrease the fSNR at the single voxel level, while the spatial covariance of the measurement noise that arises from physiological processes will remain the same. In our experience, most representation can be detected and judged at relatively coarse spatial resolutions, given the intrinsic smoothness of cortical representations (Wiestler et al., 2011), and the point-spread function of the hemodynamic response. Therefore a voxel size of 2*mm*^3^ often provides a good compromise between fSNR and spatial resolution.

Finally, our results highlight a few pitfalls that should be avoided:

- Selecting voxels or parts of the region based on the same data that is used to estimate the correlation. When selecting voxels or regions with high fSNR values (or equivalently, high split-half reliabilities or high decoding accuracies), you will overestimate the overall signal variance, and in turn underestimate the the absolute size of the correlation. If a voxel- or subregion selection is desired, it must be performed on independent functional data.
- Excluding group bootstrap estimates with fSNR estimates of zero. This will result in a bootstrap distribution that does not contain enough of the extreme values (*ρ* = 1, *ρ* = *−* 1) and Type-I error rates for testing the hypothesis of *ρ < x* will inflate dramatically. Simply retain all bootstrap estimates.

### 4.2 Current limits and possible improvements of bootstrap procedure

In this paper we used a simple bootstrap procedure, resampling subjects and calculating the percentiles of the bootstrap distributions to create confidence intervals. We then used the complement of the confidence interval as a rejection region for hypothesis testing. While this approach led to approximately correct results, the properties of the confidence interval could likely be improved by resampling the data under the Null-hypothesis (Martin, 2007; Hall and Wilson, 1991). In our case this would require the generation of artificial datasets with a specific correlation value. While our generative model (Eq. 6-8) and the MLEs of all parameters could be used to generate new artificial data under any hypothesis, the problem is that the distribution of the correlation estimate also depend on the spatial covariance of the signal and the spatial covariance of the noise. Estimating these properties reliably, such that the relevant properties of the simulated data matches those of the real data, turns out to be a hard problem.

Furthermore, the simple bootstrap does perform poorly for fewer than 20 subjects Efron et al. (1994). Being able to generate more equivalent artificial datasets would again be beneficial and would extend the applicability of the methods proposed here.

### 4.3 Application to electrophysiological and other types of data

We have focused here on the application of our methods to fMRI data. However, the problem addressed here also arises with other types of neural recording, such as electrophysiological or optical imaging data Saccenti et al. (2020). While the methods developed here can be applied in these setting, a few caveats have to be considered. First, the assumption of a Gaussian likelihood for electrophysiological measured spike rates is not appropriate, as these are closer to a Poisson distribution. A variance-stabilizing transform (such as the square root, Yu et al., 2009) is therefore a necessary step for the appropriate application of the proposed method. Furthermore, for direct electrophysiological recording, the fSNR is usually substantially higher than for fMRI, which makes the cross-block estimate (Eq. 15) a convenient shortcut for obtaining the MLE.

Of course, the estimation of the correlation of two latent (unobserved) variables is a common problem that occurs in many scientific fields (Rosner and Willett, 1988; Borrelli and Cole, 1990; Beaton et al., 1979; Liu et al., 1978). While each setting has its own specific challenges, the maximum-likelihood approach using a mixture of fixed and random effects (Eq. 18) can be flexibly adapted to match the data characteristics.

### 4.4 Cosine similarities and representational geometry

We presented here a methods that allows us to test predictions about the correlation or angle between two high-dimensional vectors that are measured with noise (Fig. 1b,c). In neuroscience, this provides an important tool for testing a large range of hypotheses about representational geometries (Kriegeskorte and Wei, 2021). To illustrate the breadth of hypotheses that can be tested, we discuss two examples.

First, take the example of an animal or human learning to make a response to two items (a,b) under two conditions (red or blue, Fig. 10a-c). In this setting, the geometry of the representation provides important insight about how a biological systems learn rules and is then able to apply these in a flexible manner (Bernardi et al., 2020; Xie et al., 2022; Boyle et al., 2024). If the four stimuli *times* condition combinations are represented, such that all difference between the combinations are orthogonal (*ρ* = 0), the representation would allow the separation of any pairs of items (Fig. 10a). Thus, such a representation would allow for learning any possible decision rule. The disadvantage, however, is that a rule learned under condition X would not generalize to condition Y. In contrast, in a representation in which the difference between the two stimuli is parallel across conditions (*ρ* = 1, Fig. 10b), a rule learned in one condition would generalize fully to the other. This representation would not allow learning of any rule, however. For example the XOR problem of separating **x**_*a*_, **y**_*b*_ from **x**_*b*_, **y**_*a*_ would not possible using a linear readout. An intermediate representation, in which the two item differences are not entirely parallel, but slightly tilted along an orthogonal axis (*ρ <* 1, Fig. 10c), unifies both advantages, having both the ability to generalize and to learn new rules along any other dimension (Bernardi et al., 2020). In this scenario, the exact angle between two lines connecting different pairs of items can serve as an important measure of the balance between abstractness and flexibility of the representation.

**Figure 10.**
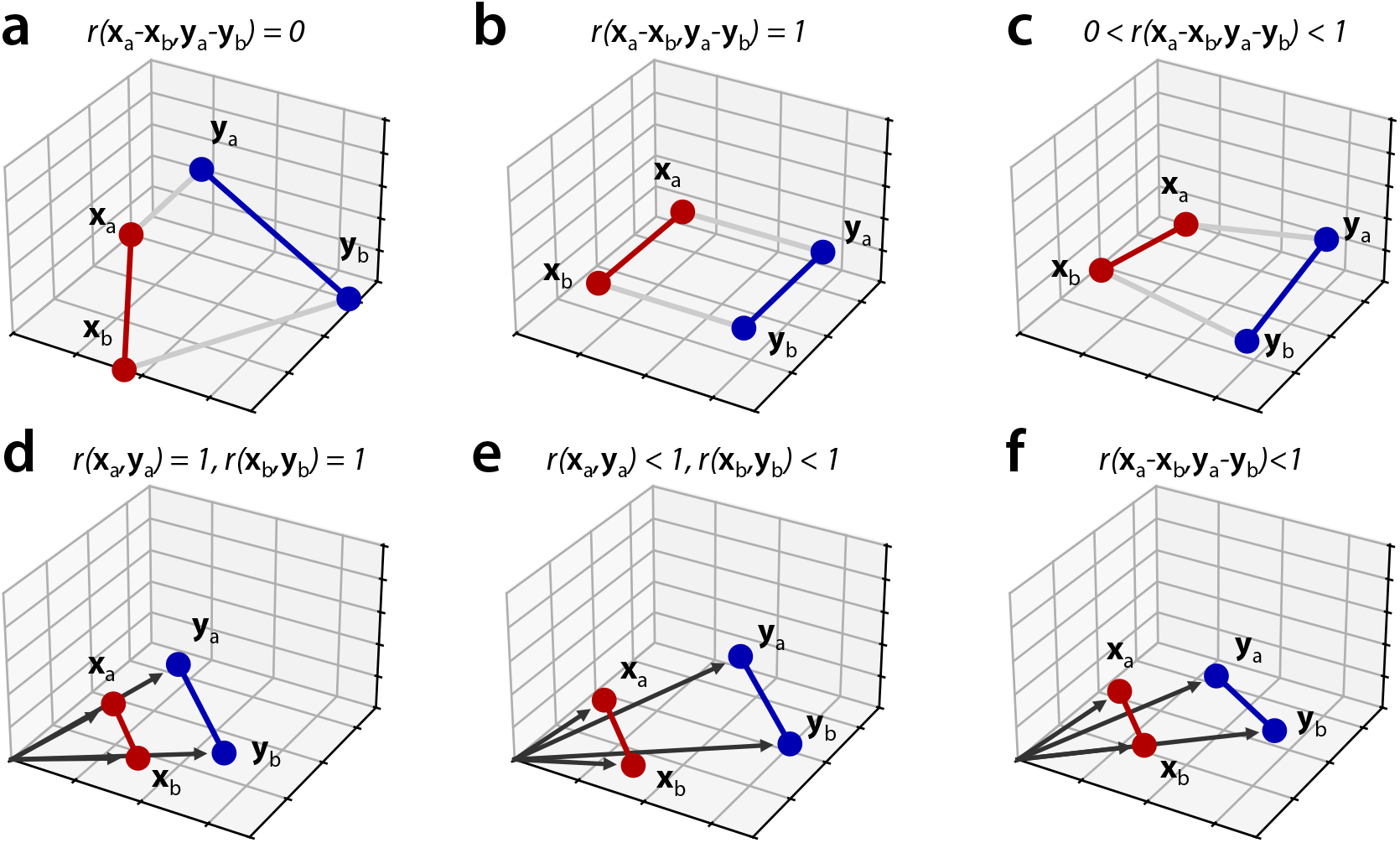
Common hypotheses representational geometries that can be tested using cosine similarities. **Top row:** How are items (a,b) are represented under two conditions (x,y)? **(a)** Independent representation allow full flexibility in learning, but no generalization. **(b)** Perfectly aligned representations allows for generalization of knowledge learned in one condition to generalize to the other condition, **(c)** Partially aligned representations allow for both generalization and condition-specific learning, with the angle of alignment characterizing the precise solution. **Bottom row:** How do activation patterns for two items (a,b) scale relative to rest (origin) under two conditions (x,y)? **(d)** The activation pattern for each item increases proportionally relative to rest. **(e)** Additional to an increase in activity, there is a additive shift, causing imperfect scaling. However the difference vector between item a and b remains parallel across conditions. **(f)** Across conditions, activity increases, but also leads to a item-specific change, such that the difference vectors are not perfectly aligned.

Another commonly occurring example are the changes in the patterns of two or more items, when the overall activation increases (Fig. 10d-f). One example here is finger representations in M1 or S1 when the speed (Arbuckle et al., 2019) or force (Diedrichsen et al., 2013) of the movement increases. Similar problems occur when we study the effects of repetition suppression onto representations (Berlot et al., 2021). In these cases, we may want test the hypothesis of pure scaling, which predicts that the vector **x**_*a*_ is perfectly parallel to **y**_*a*_, as well as **x**_*b*_ to **y**_*b*_. We may also have a situation in which each finger-specific pattern scales, but an additional overall background pattern also increases (Arbuckle et al., 2019). In this case, we would predict that the above cosine similarities would be smaller than 1, but that the vector between the two fingers **x**_*a*_*−* **x**_*b*_ be parallel to **y**_*a*_ *−***y**_*b*_. If our inference suggests that the cosine similarity between these two vectors is smaller than one, it would provide evidence that the representation of finger movements in M1 changes at higher activation levels - with 1-cosine similarity quantifying the strength of the change. This would indicate that the brain does not simply engage the the same system more for faster or stronger actions, but qualitatively changes the control.

These examples, hopefully, illustrate how complex and interesting questions about the representational geometry often can be translated into simple hypotheses about the size of the real cosine similarity (or correlation) between pattern vectors. We hope therefore that the methods tested here provide a useful tool to test these such hypothesis on fMRI and neural data.

## 5 ENDING SECTIONS

### Code Availability

The maximum-likelihood estimation is implemented in the PcmPy toolbox (version 1.2), available at https://github.com/DiedrichsenLab/PcmPy. An tutorial of how to use the toolbox to implement the methods used in this paper can be found at https://pcm-toolbox-python.readthedocs.io/en/latest/examples.html.

### Author Contributions

Conception: JD, SB; Implementation: JD; Simulations: JD, XF; Empirical data application: MS, JD; Draft: JD, SB; Final editing: JD, XF, MS, SB.

### Funding

This study was supported by Discovery Grants from the Natural Sciences and Engineering Research Council of Canada (NSERC, RGPIN 2016-04890 to JD and RGPIN 2023-03894 to SB.). Additional funding came from the Canada First Research Excellence Fund (BrainsCAN) to Western University. XF was supported through an Undergraduate Student Research Internship award from the University of Western Ontario.

## Acknowledgments

The authors thank Nikolaus Kriegeskorte, Giacomo Ariani, Eva Berlot, and Marco Emanuele for useful discussion and feedback.

## APPENDICES

### 6.1 Symbol table

**Table 1.** Meaning of mathematic symbols used in the paper for the single-pattern setting.

| Symbol | Size | Meaning |
| --- | --- | --- |
| $x_p^*$ | | True activation for condition $X$ in voxel $p$ |
| $y_p^*$ | | True activation for condition $Y$ in voxel $p$ |
| $x_{i,p}$ | | $i^{th}$ measure of condition $X$ in voxel $p$ |
| $y_{i,p}$ | | $i^{th}$ measure of condition $Y$ in voxel $p$ |
| $P$ | | Number of voxels |
| $n_x$ | | Number of measurement for $X$ |
| $n_y$ | | Number of measurements for $Y$ |
| $N$ | | Total number of measurement |
| $\mathbf{x}^*$ | $1 \times P$ | True activation pattern for condition $X$ |
| $\mathbf{y}^*$ | $1 \times P$ | True activation pattern for condition $Y$ |
| $\mathbf{x}_i$ | $1 \times P$ | $i^{th}$ measure of pattern for condition $X$ |
| $\mathbf{y}_i$ | $1 \times P$ | $i^{th}$ measure of pattern for condition $Y$ |
| $\rho$ | | correlation of true patterns across voxels |
| $\sigma_\epsilon^2$ | | variance of measurement noise |
| $\sigma_x^2$ | | variance of $x^*$ across voxels |
| $\sigma_y^2$ | | variance of $y^*$ across voxels |
| $\mathbf{d}_p$ | $N \times 1$ | All observations for a single voxel |
| $\mathbf{D}$ | $N \times P$ | Entire dataset for a single subject |
| $\mathbf{u}_p$ | $2 \times 1$ | True patterns for voxel $p$ |
| $\mathbf{G}$ | $2 \times 2$ | Covariance matrix for true patterns |
| $\mathbf{Z}$ | $N \times 2$ | Design matrix linking observations to conditions |
| $\mathbf{V}$ | $N \times N$ | Covariance matrix for observations $\mathbf{d}$ |

**Table 2.** Meaning of additional mathematic symbols used in the paper for the multi-pattern setting.

| Symbol | Size | Meaning |
| --- | --- | --- |
| $K$ | | Number of items |
| $x_{j,p}^*$ | | True activation for $j^{th}$ item in condition $X$ in voxel $p$ |
| $y_{j,p}^*$ | | True activation for $j^{th}$ item in condition $Y$ in voxel $p$ |
| $x_{i,j,p}$ | | $i^{th}$ measure for $j^{th}$ item in condition $X$ in voxel $p$ |
| $y_{i,j,p}$ | | $i^{th}$ measure for $j^{th}$ item in condition $Y$ in voxel $p$ |
| $\tilde{x}_{i,j,p}$ | | Same as above, but mean across items removed |
| $\tilde{y}_{i,j,p}$ | | Same as above, but mean across items removed |
| $\mathbf{u}_p$ | $2K \times 1$ | Item-specific deviations from condition mean for voxel $p$ |
| $\mathbf{m}_p$ | $2 \times 1$ | Condition means for voxel $p$ |
| $\mathbf{G}$ | $2K \times 2K$ | Covariance matrix of item-specific patterns |
| $\mathbf{Z}$ | $N \times 2K$ | Design matrix linking observations to conditions and items |
| $\mathbf{X}$ | $N \times 2$ | Design matrix linking observations to conditions |
| $\mathbf{V}$ | $N \times N$ | Covariance matrix for observations $\mathbf{d}$ |

### 6.2 Cross-block estimates with separate noise covariances

If we assume separate measurement noise variances 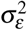 and 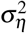 for *x* and *y*, respectively), the cross-block estimators of our variances become:

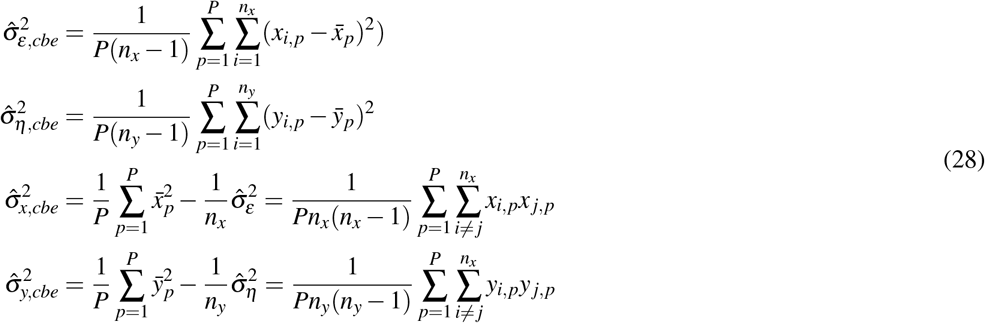

from the last two equations, we can see that the signal variance is estimated by the covariance of the patterns across different runs (hence the name cross-block estimator). The correlation then can be estimated as before using equation 15.

### 6.3 Optimization details

To optimize the log-likelihood in Eq. 9, we use a Newton-Raphson algorithm (Lindstrom and Bates, 1988) with the expected second derivative, the Fisher-information matrix:

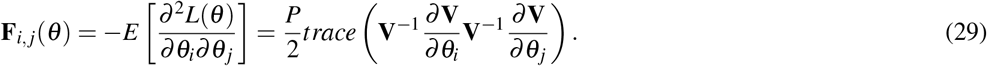

### 6.4 Restricted Maximum Likelihood

To estimate the parameters *θ*) in the presence of fixed effects **X**, we defined the the residual forming matrix:

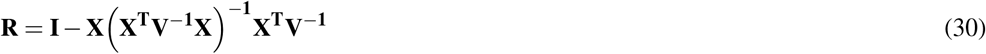

The Restricted log-likelihood function can then be written as:

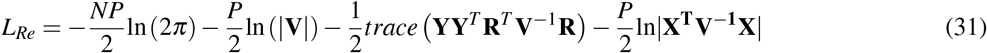

Note that the third term can be simplified by noting that

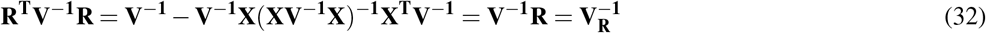

The derivative of each parameter then is:

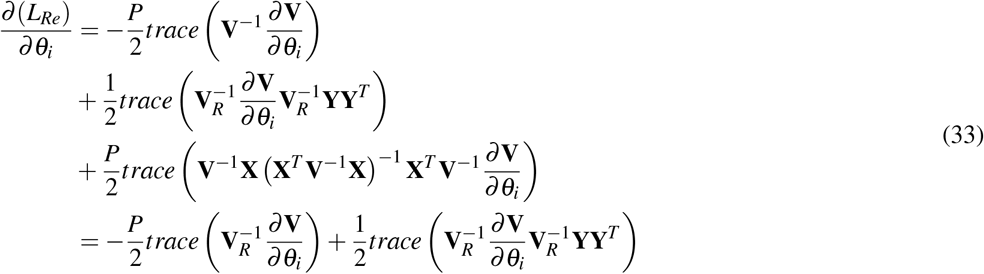

The Fisher-information matrix then is:

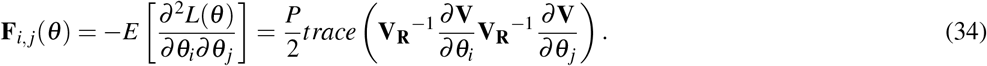

### 6.5 Unequal fSNR across the two conditions

We repeated the simulations from Fig. 2, using unequal fSNRs for the two conditions. We set *ρ* = 0.7, 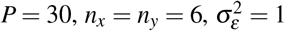 1, and varied log 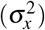 and log 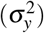 between *−*6 and 2. The mean (Fig. 11a) and the mean absolute error (Fig. 11b) of the maximum-likelihood estimates shows the dependence on both signal variances. For a log-variances difference for up to 2, the MLE behaved similar to simulation in which the two signal variance were equal (for the same mean log signal variance). For larger differences, the behavior of the MLE was more influenced by the lower log-signal variance. A difference in log-variance of 2 is equivalent to a ratio of fSNRs of *exp*(2) = 7.389.

**Figure 11.**
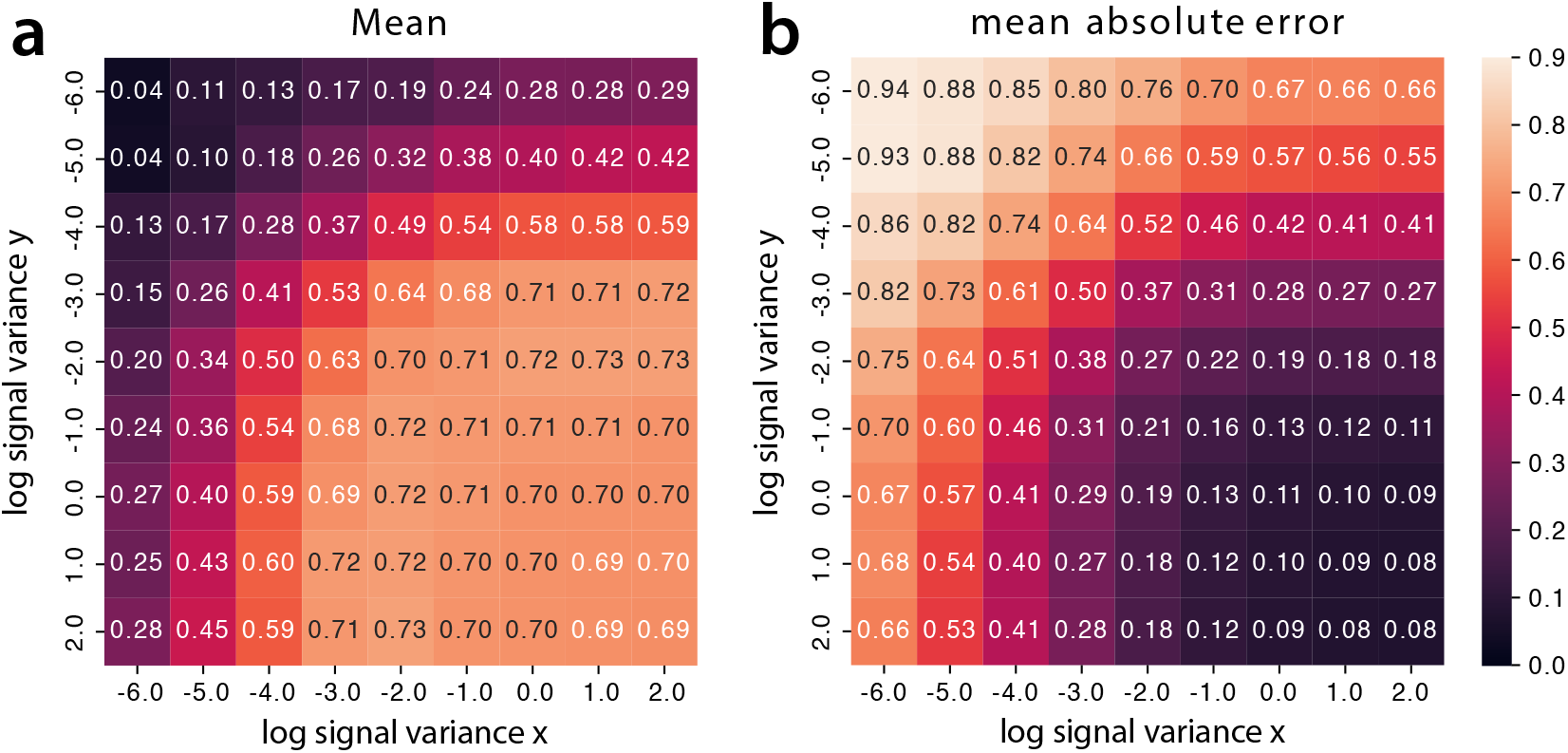
MLE estimate for unequal signal variances across conditions (x,y). **(a)** Mean maximum-likelihood correlation estimate (no exclusion) as a function of log 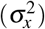 and log 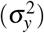. The true value is *ρ* = 0.7. **(b)** Mean absolute error of correlation estimate as a function of the two signal variances. Note that 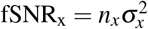.

## Footnotes

1 Note that we are using the term correlation to include both the Pearson correlation (in which the mean value across voxels in each condition is subtracted), as well as the cosine similarity (in which the mean value across voxels is not subtracted, see methods). The results presented in the paper pertain to both situations.

